# Behavioral Efficacy of AAV FOXG1 Gene Replacement Therapy in a Mouse Model of FOXG1 Syndrome

**DOI:** 10.1101/2025.04.02.646887

**Authors:** Christina L. Torturo, Ailing Du, Kimberly Kerker, Jodi Gresack, Christopher DeGrave, Kathleen Funk, Princy Agnihotri, Frank Hsieh, Teresa M. Gunn, Sylvie Ramboz, Brian Bettencourt, Scott Reich, Guangping Gao, Dan Wang, Dinah W. Y. Sah

## Abstract

FOXG1 syndrome is a severe neurodevelopmental disorder characterized by microcephaly, profound intellectual disability with communication deficits including lack of speech, impaired social interaction, increased anxiety, hyperkinetic/dyskinetic movements, seizures and abnormal sleep patterns. Mutations in a single allele of the *FOXG1* gene cause disease, likely due to loss-of-function. However, current therapies do not target this root cause of FOXG1 syndrome and have little to modest therapeutic benefit on only a small subset of symptoms. To date, the therapeutic potential of restoring FOXG1 levels in the brain with adeno-associated virus (AAV) *FOXG1* gene replacement therapy has only been reported in a *Foxg1fl/+;NexCre* mouse model that lacks one *Foxg1* allele but does not express mutant FOXG1, and with only neuroanatomical endpoints evaluated. Here, in a FOXG1 mouse model that contains a highly prevalent, patient-specific Q84P mutation, we describe the beneficial effects of AAV human *FOXG1* gene replacement therapy administered by intracerebroventricular (ICV) injection at postnatal day 6 (P6) on several behavioral deficits that are relevant to key features of human FOXG1 syndrome. Our studies demonstrate that AAV *FOXG1* gene replacement therapy is a promising approach for the treatment of a subset of functional deficits in human FOXG1 syndrome.

## INTRODUCTION

FOXG1 syndrome is a severe neurodevelopmental disorder caused by mutations in a single allele of the *FOXG1* gene. Clinical manifestations of FOXG1 syndrome, which is an autism spectrum disorder, include severe global developmental delay, postnatal growth deficiency, microcephaly, profound intellectual disability with communication deficits including lack of speech, impaired social interaction, increased anxiety, hyperkinetic/dyskinetic movements, seizures, abnormal sleep patterns, and gastrointestinal and feeding problems (Wong *et al.* 2019). Onset of most symptoms typically occurs within the first year of life with most diagnoses between birth and 3 years of age (Brimble *et al.* 2023). The clinical manifestations have a profound deleterious impact on the patient’s quality of life. Current therapies are symptomatic and have little to modest therapeutic benefit on only a small subset of symptoms. Thus, there is a high unmet medical need for effective treatments that target the root cause of disease.

FOXG1 is a forkhead family transcription factor that is critical for forebrain development and territorial specification, including neurogenesis and balance of excitatory versus inhibitory neuronal differentiation, neurite outgrowth, organization of the cortex, lamination and cortico-cortical connections (Wong *et al.* 2019). In addition, FOXG1 plays a key role in dendritogenesis and neural plasticity (Wong *et al.* 2019). Mutations in the *FOXG1* gene that cause FOXG1 syndrome include frameshift, missense, nonsense and deletion variants that result in loss or gain of function (Frisari *et al.* 2022, Brimble *et al.* 2023). The decrease in levels of functional FOXG1 protein disrupts neurodevelopment, leading to abnormalities that include reduced cortical layers and numbers of cortical neurons, atrophy of the hippocampus, agenesis of the corpus callosum and delay in myelination (Pringsheim *et al.* 2019) that contribute to the clinical manifestations.

Human FOXG1 syndrome is believed to be caused by haploinsufficiency. In a mouse model of FOXG1 haploinsufficiency, deletion of one *Foxg1* allele in cortical, hippocampal and hypothalamic neurons and in intermediate progenitor cells (*Foxg1fl/+;NexCre* or *Foxg1*-cHet; Jeon *et al.* 2024) resulted in microcephaly, corpus callosum malformation, reduction of the cortical upper layers, and hippocampus dentate gyrus malformation, similar to structural changes in the human disease. In this mouse model, ICV administration of adeno-associated virus 9 (AAV9) with a human FOXG1 payload driven by a chicken beta-actin (CBA) promoter (AAV9.CBA.hFOXG1) on postnatal day 1 was reported to result in human FOXG1 expression throughout the brain, not only in cells that normally express FOXG1 but also in those that do not normally express FOXG1 (Jeon *et al.* 2024). In conjunction with this expression of FOXG1, some but not all neuroanatomical deficits were ameliorated. The authors reported nearly complete normalization of (A) the increased number of oligodendrocyte progenitor cells (OLIG2+) in the corpus callosum, (B) the reduced number of CTIP2+ cortical neurons (possibly by preventing neuronal cell death), and (C) the shortened length and widened apex angle of dentate gyrus (dentate gyrus abnormalities). Partial normalization of both reduced thickness of the corpus callosum, and reduced myelin basic protein-immunoreactivity in the corpus callosum, reflecting myelination deficits, was also observed. However, there was no effect of AAV9.CBA.hFOXG1 administration on the thickness of the cortex. An important consideration for interpreting results from this *Foxg1fl/+;NexCre* mouse model is that in contrast to human disease, mutant FOXG1 is not present in cortical, hippocampal and hypothalamic neurons or in intermediate progenitor cells, which may impact the phenotype and/or ability to ameliorate the phenotype with AAV9 *FOXG1* gene replacement therapy. Furthermore, it is unclear how the improvement in some but not all neuroanatomical deficits impacts dysfunction, including behavioral deficits.

To date, there have been no reports published on the behavioral effects of AAV *FOXG1* gene replacement therapy in a mouse model of FOXG1 syndrome. To conduct these studies, we used a newly developed C57BL/6J-Foxg1^em1Mri^ mouse model (C57BL/6J-Foxg1^Q84P^, which carries a duplication of a cytosine at position 250 of the coding sequence, c.250dupC, resulting in p.Q84Pfs30X) courtesy of Dr. Teresa Gunn, McLaughlin Research Institute, that contains a frameshift mutation in one allele of *Foxg1*, encoding a mutant, truncated form of murine FOXG1 protein. This Q84P mouse model more closely represents the genotype of a prevalent form of human FOXG1 syndrome (p.Q86Pfs35X) since a mutant *Foxg1* gene is present, unlike the *Foxg1fl/+;NexCre* mouse model which does not express mutant FOXG1.

Here, we describe studies in this Q84P mouse model to characterize behavioral phenotype and evaluate the effects of AAV human *FOXG1* gene replacement therapy administered by ICV injection at P6 on behavioral deficits. To administer the AAV *FOXG1* gene replacement therapy in heterozygous Q84P mice, we selected the ICV route that has been used for delivering therapeutic levels of payload expression to the central nervous system for multiple neurological disorders. ICV administration provides substantially increased efficiency of delivery to the brain while reducing distribution to peripheral tissues and organs. Importantly, for clinical translation, ICV dosing avoids neutralizing antibodies to the capsid that may be present in circulation. We designed the components of the AAV vector to: (A) maximize distribution to cortical and hippocampal neurons - critical cells for the expression of FOXG1 to treat FOXG1 syndrome, and (B) provide the shortest time to onset of transgene expression for attaining significant therapeutic benefit as quickly as possible. We selected the AAV9 capsid because it has been administered in the clinic and provides widespread brain distribution after ICV injection.

A self-complementary (sc) rather than single-stranded payload was used for the most rapid onset of *FOXG1* transgene expression, and the human *FOXG1* transgene was codon-optimized for enhanced expression. Given the predominant role of FOXG1 expression in neurons, we chose the neuronal human synapsin-1 (hSyn1) promoter. Thus, we evaluated scAAV9.hSyn1-opthFOXG1 administered by ICV injection in the Q84P mouse model of FOXG1 syndrome. Furthermore, we focused on postnatal day 6 (P6) for AAV administration based on translational considerations of balancing disease treatment as early as possible with clinical practice given time to diagnosis.

Vector genome (VG), human *FOXG1* mRNA and human FOXG1 protein levels were quantified to confirm AAV delivery and expression of exogenous human FOXG1 in the brain. To characterize the behavioral phenotype in this mouse model, we evaluated ultrasonic vocalization at postnatal day 9 (P9), and open field, clasping, grip strength, wire hang, tapered balance beam, running wheel, acoustic startle, fear conditioning, Y-maze, and SmartCube^®^ endpoints at 4 to 13 weeks of age. There were significant deficits in ultrasonic vocalization at P9, open field, running wheel, grip strength, contextual fear conditioning, Y-maze exploratory activity and a subset of SmartCube^®^ endpoints compared to wild-type mice, but no differences in clasping, wire hang, tapered balance beam, acoustic startle or cued fear conditioning. The grip strength phenotype was mild and observed in only one of two studies. ICV administration of scAAV9.hSyn1-opthFOXG1 at P6 resulted in dose-related improvements in a number of open field, running wheel and SmartCube^®^ deficits, but not fear conditioning endpoints. Ultrasonic vocalization at P9 was not evaluated because administration of AAV at P6 would not provide sufficient time for transgene expression. All tested ICV doses of scAAV9.hSyn1-opthFOXG1, including the efficacious top dose, were well-tolerated with no significant effects on body weight, no abnormal cage side observations and no microscopic abnormalities in brain structure.

These results in the Q84P mouse model of FOXG1 syndrome demonstrate that ICV administration of AAV *FOXG1* gene replacement therapy comprising scAAV9.hSyn1-opthFOXG1, with consequent increased levels of human FOXG1 in critical brain regions, ameliorates several but not all behavioral deficits that are relevant to key features of human FOXG1 syndrome and is well-tolerated at efficacious doses. Importantly, administration at P6 was effective, suggesting that ICV treatment at later childhood ages, which is potentially more practical for clinical translation than treatment at birth, may be beneficial. Our studies indicate that ICV administration of scAAV9.hSyn1-opthFOXG1 is a promising approach to the treatment of some aspects of human FOXG1 syndrome.

## RESULTS

To evaluate the behavioral effects of AAV *FOXG1* gene replacement therapy, we used a newly developed Q84P mouse model of FOXG1 syndrome. This Q84P mouse model is heterozygous on a C57BL/6 background and was created with CRISPR editing to mutate the 84th codon by inserting a guanine nucleotide into the noncoding (antisense) strand of the murine *Foxg1* locus, resulting in a complementary cytosine nucleotide inserted into the coding (sense) sequence. This mutation changed the encoded amino acid from glutamine to proline and also shifted the frame of the coding sequence to introduce a premature stop codon after an additional 44 codons. The resultant murine FOXG1 protein is both mutated and truncated to 113 amino acids in length, compared to the wildtype murine FOXG1 protein which is 481 amino acids in length.

### Phenotypic characterization of the Q84P mouse model reveals body weight, ultrasonic vocalization, open field, Y-maze, fear conditioning and grip strength phenotypes

To identify behavioral endpoints with substantial differences between Q84P and wild-type (WT) mice that could be affected by AAV *FOXG1* gene replacement therapy, we began by characterizing the baseline phenotype of the Q84P mouse model.

#### Survival, body weight, and neonatal well-being, motor and ultrasonic vocalization tests

There was no difference in survival between Q84P and WT mice evaluated from birth to 12 weeks of age. All animals survived with the exception of one WT female, one WT male and 2 Q84P males that died after submandibular blood collections at 7 weeks of age, likely related to the procedure.

Body weights (males and females combined) showed a significant transient deficit in Q84P compared with WT animals on postnatal day 20 (P20; **Fig. 1A**), and at 3, 4, 5, and 6 weeks of age (**Fig. 1B**). Male Q84P mice exhibited significantly lower body weights than male WT mice at 4, 5, 6, 7, 8 and 12 weeks of age, whereas female Q84P mice had significantly lower body weights than female WT mice at only 4 and 5 weeks of age (**SUPPLEMENTARY Fig. 1**). These results indicate that the Q84P body weight phenotype is present transiently in young adult mice.

**Figure 1.**
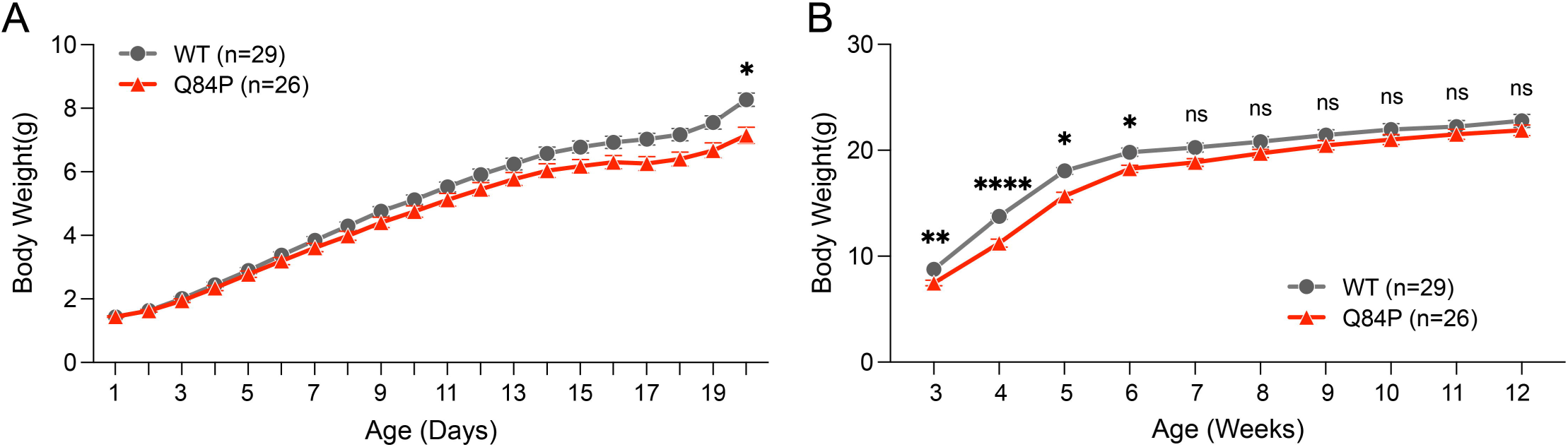
Body weights of Q84P mice from postnatal day 1 to week 12. (**A**) Male and female neonatal body weights combined showed significant differences between the genotypes as assessed by one-way ANOVA (F (1, 53) = 4.069, p = 0.0488), with a significant post-hoc difference only at postnatal day 20 (P20), as assessed by Tukey’s multiple comparisons test. (**B**) Male and female adult body weights combined showed significant differences between the genotypes as assessed by one-way ANOVA (F (1, 53) = 5.447, p = 0.0234), with significant post-hoc differences at weeks 3, 4, 5 and 6, as assessed by Tukey’s multiple comparisons test. The number of animals per group (n) reflect the total number of animals in each group at the time of enrollment. Means ± standard error of the mean are displayed. ns: not significant. * p < 0.05, ** p < 0.01, **** p < 0.0001

Neonatal well-being indices (gasping, skin color, milk content and muscle tone), assessed every 48 hours from birth through postnatal day 16 (P16), showed no phenotypic differences between Q84P and WT mice. Respiration rate and body temperature assessed every 4 days until P20 also showed no phenotypic differences between Q84P and WT mice.

Neonatal motor tests including the righting reflex (**SUPPLEMENTARY Fig. 2A**), geotaxis (**SUPPLEMENTARY Fig. 2B**) and tube (**SUPPLEMENTARY Fig. 2C**) tests, evaluated through postnatal day 14 (P14), showed no consistent phenotypic differences.

Neonatal ultrasonic vocalization (USV) measurements at P9 (**SUPPLEMENTARY Fig. 2D**) revealed that Q84P mice vocalized significantly less than WT mice (males and females combined) which was driven by a significant phenotype in female Q84P mice vs WT mice. The reduction in USV suggests that P9 female Q84P mice have a communication deficit.

#### Open field

Q84P mice exhibited a significant open field phenotype compared to WT mice at all 3 ages evaluated (4, 6 and 8 weeks of age). Q84P mice travelled significantly less total (**Fig. 2A**) and center (**Fig. 2B**) distance than WT mice (males and females combined). Both male and female Q84P mice demonstrated these phenotypes.

**Figure 2.**
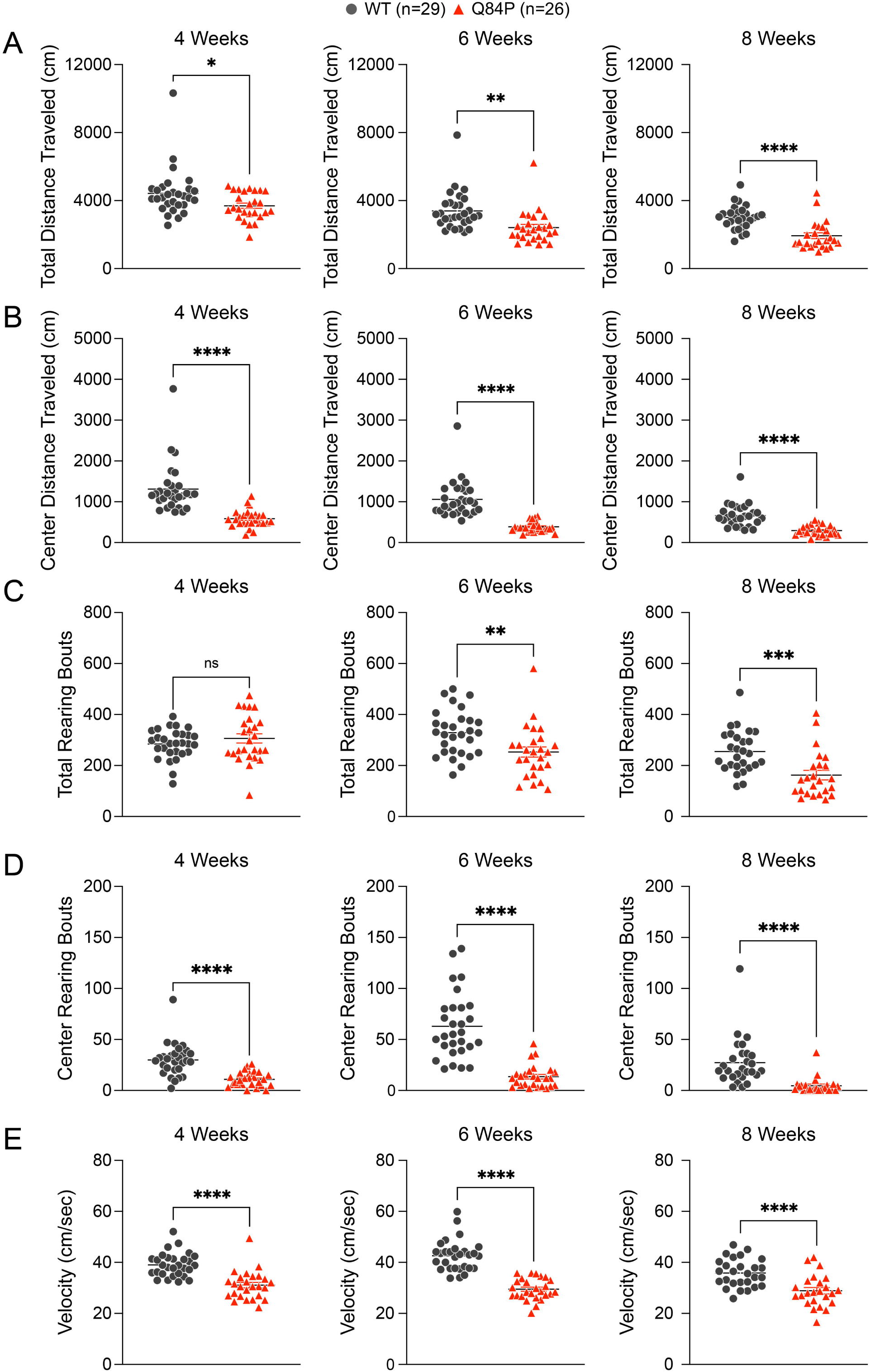
Open field distance travelled, rearing frequency and velocity show significant reductions in Q84P mice compared to WT mice, with male and female animals combined at 4, 6 and 8 weeks of age. Open field total (**A**) and center (**B**) distance travelled, center rearing frequency (**D**) and velocity (**E**) showed significant differences between genotypes at 4, 6 and 8 weeks of age. Open field total rearing frequency (**C**) showed significant differences between genotypes at 6 and 8 but not 4 weeks of age. Statistical comparisons were assessed by unpaired t-test. The number of animals per group (n) reflect the total number of animals in each group at the time of enrollment. Means ± standard error of the mean are displayed. ns: not significant. * p < 0.05, ** p < 0.01, *** p < 0.001, **** p < 0.0001

Q84P mice also exhibited less frequent total (**Fig. 2C**) and center (**Fig. 2D**) rearing, and decreased open field velocity (**Fig. 2E**) at all ages evaluated except for total rearing at 4 weeks of age. Total rearing frequency (males and females combined) was reduced in Q84P compared with WT mice at 6 and 8 weeks of age, which was driven by a significant phenotype in female Q84P mice. Center rearing frequency was also significantly lower in Q84P compared with WT mice (males and females combined at 4, 6 and 8 weeks of age), with both males and females demonstrating this phenotype, though more pronounced in females. Open field velocity at 4, 6 and 8 weeks of age was reduced significantly in Q84P versus WT mice (males and females combined), with both males and females demonstrating this phenotype.

Taken together, these open field deficits of reduced activity (less total and center distance travelled, less total and center rearing frequency, and lower velocity) in Q84P mice at 4, 6 and/or 8 weeks of age suggest that they have higher levels of anxiety and/or deficits in exploratory behavior compared to WT mice.

#### Y-maze

Exploratory behavior and working memory were evaluated in the Y-maze at 9 weeks of age by quantifying total number of arm entries (**Fig. 3A**) and arm alteration (**Fig. 3B**), respectively. There was a significant deficit in the total number of Y-maze arm entries in Q84P compared with WT mice (males and females combined) which was driven by a significant phenotype in the female Q84P mice versus female WT mice (**SUPPLEMENTARY Fig. 3A**). There were no phenotypic differences in percent alteration. These results suggest that Q84P mice have a deficit in exploratory activity but no deficit in working memory.

**Figure 3.**
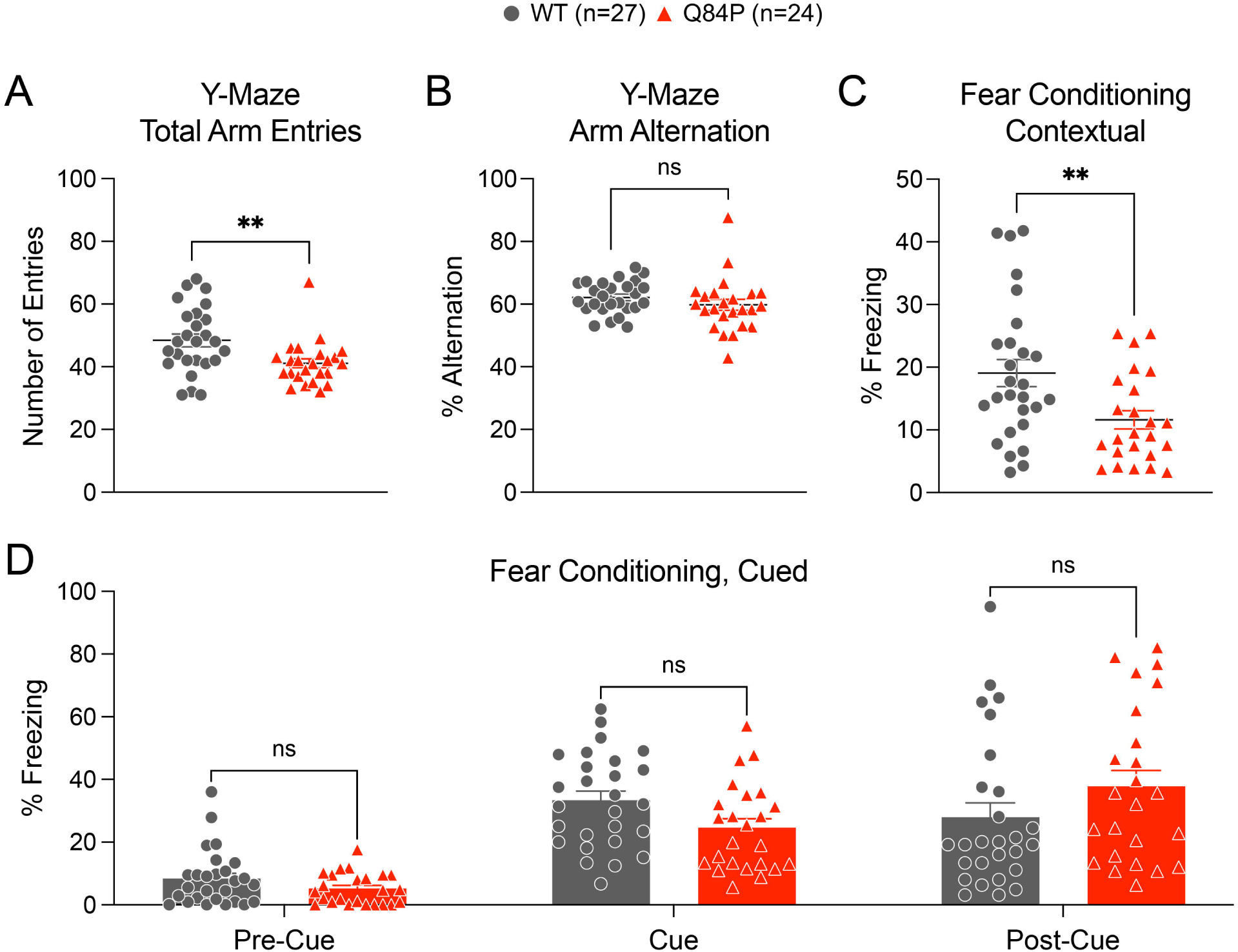
Y-maze total arm entries and contextual fear conditioning are reduced in Q84P mice compared to WT mice, with male and female animals combined. Y-maze total arm entries (**A**) but not Y-maze percent arm alteration (**B**) showed a significant difference between the genotypes at 9 weeks of age, as assessed by unpaired t-test. (**C**) Contextual fear conditioning showed a significant difference between the genotypes at 10 weeks of age as assessed by unpaired t-test. (**D**) Cued fear conditioning was not significantly different between the genotypes in male and female mice combined, as assessed by two-way ANOVA with Sidak’s multiple comparisons post-hoc test (F(1, 49) = 0.4812, p = 0.8273). Means ± standard error of the mean are displayed. ns: not significant. ** p < 0.01

#### Fear conditioning

Contextual (**Fig. 3C**) and cued (**Fig. 3D**) fear conditioning, assessed at 10 weeks of age, showed that Q84P mice exhibited significantly less freezing in contextual fear conditioning than WT mice (males and females combined) which was driven by a significant phenotype in female Q84P mice versus female WT mice (**SUPPLEMENTARY Fig. 3B**). Neither female nor male Q84P mice exhibited a phenotype in freezing in cued fear conditioning (**SUPPLEMENTARY Fig. 3C**).

#### Acoustic startle

The acoustic startle response at 6 and 8 weeks of age showed no phenotypic differences between Q84P and WT mice (**SUPPLEMENTARY Fig. 4**).

#### Grip strength

Q84P mice exhibited a significant reduction in grip strength compared to WT mice at 4, 6, and 8 weeks of age, with these differences declining across these ages and becoming insignificant by 10 and 12 weeks of age (**Fig. 4**). Male Q84P mice exhibited significant grip strength deficits at 4, 6 and 8 weeks of age whereas female Q84P mice showed significant deficits only at 4 weeks of age (**SUPPLEMENTARY Fig. 5**). These results suggest that the grip strength phenotype is present transiently in young adult Q84P mice and more pronounced in males than in females.

**Figure 4.**
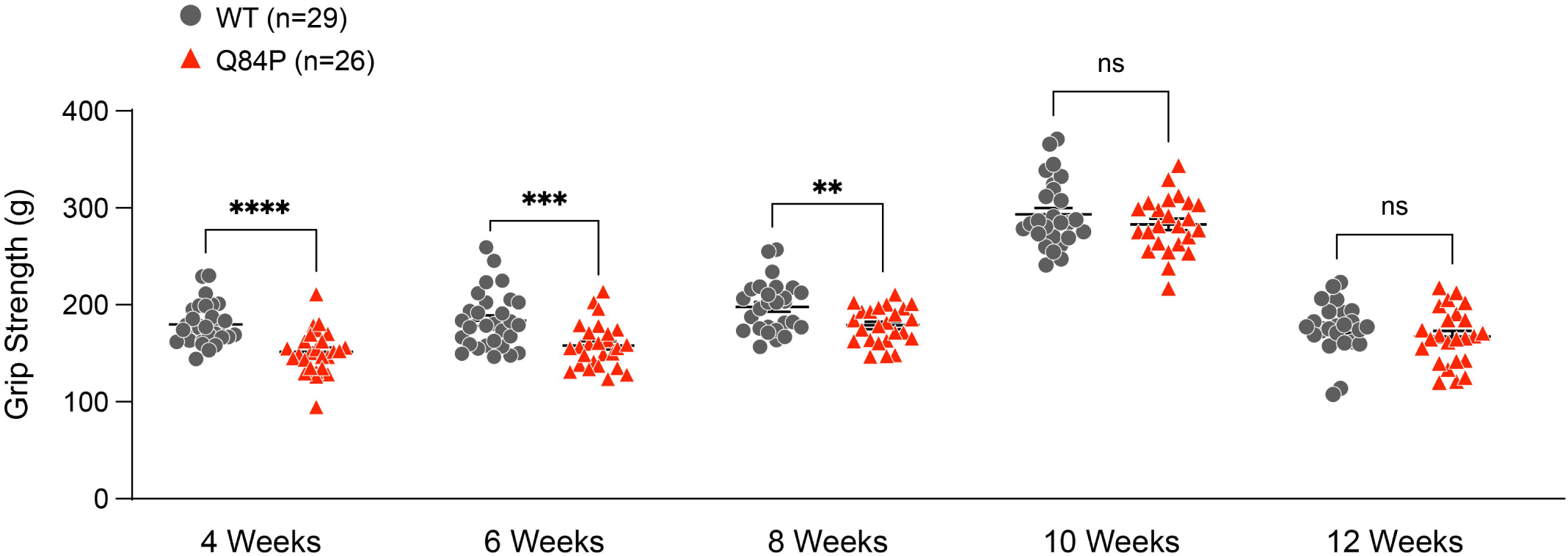
Grip strength is transiently reduced at 4, 6 and 8 weeks of age in Q84P mice compared with WT mice. Grip strength in males and females combined, averaged over 5 trials, showed significant differences between genotypes at 4, 6 and 8 but not 10 or 12 weeks of age. Statistical comparisons were assessed by unpaired t-test. The number of animals per group (n) reflect the total number of animals in each group at the time of enrollment. Means ± standard error of the mean are displayed. ns: not significant. ** p < 0.01, *** p < 0.001, **** p < 0.0001

#### Clasping

Clasping behavior (**SUPPLEMENTARY Fig. 6**) showed no phenotypic differences between Q84P and WT mice at 4 or 6 weeks of age. No clasping was observed in any Q84P or WT mice at 8, 10, 11 and 12 weeks of age.

#### Neurofilament-light chain protein levels

To evaluate a biomarker of neurodegeneration, we quantified neurofilament-light (NfL) chain protein levels in cerebrospinal fluid (CSF) and plasma at 12 weeks of age (**SUPPLEMENTARY Fig. 7A** and **B**, respectively). There were no significant differences between Q84P and WT mice in CSF or plasma NfL levels, suggesting that there is no significant neurodegeneration in the Q84P mice and that functional deficits at this age are driven by neuronal dysfunction rather than neurodegeneration or axonal loss.

Taken together, in adult male and female Q84P mice, the most robust behavioral phenotypes of those characterized are open field deficits and a transient grip strength reduction. In addition, in adult female Q84P mice, Y-maze exploratory behavior and contextual fear conditioning represent significant behavioral phenotypes whereas in neonatal female Q84P mice, USV deficits are pronounced and significant.

### ICV injection of scAAV9.hSyn1-opthFOXG1 in the Q84P mouse on postnatal day 6 is well-tolerated

Two AAV studies were conducted in Q84P mice to evaluate the tolerability of AAV *FOXG1* gene replacement therapy and efficacy on behavioral phenotypes. In both studies, we administered scAAV9.hSyn1-opthFOXG1 by ICV injection in heterozygous Q84P male mice at P6. In the first study, we evaluated a range of doses of scAAV9.hSyn1-opthFOXG1 (4 × 10^10^, 1 × 10^11^ and 2.6 × 10^11^ vg per mouse) in male Q84P mice for effects on open field, fear conditioning and grip strength behaviors compared to vehicle-treated Q84P and WT mice. In a subsequent study, to confirm and extend the initial efficacy observations, we administered a single ICV dose level of 2.6 × 10^11^ vg scAAV9.hSyn1-opthFOXG1 per mouse to male Q84P mice and assessed open field, wire hang, tapered balance beam, running wheel and SmartCube^®^ behavioral endpoints. Vector genome and human *FOXG1* mRNA and human FOXG1 protein levels, as well as immunohistochemical staining for FOXG1 in the brain, were assessed to confirm delivery to key brain regions and to correlate expression levels with changes in behavioral endpoints. In addition, survival, body weight and cage side observations were used to monitor general tolerability while neuropathology status was assessed by hematoxylin and eosin staining of the brain.

#### Survival, cage side observations and neuropathology

ICV treatment of heterozygous Q84P mice with scAAV9.hSyn1-opthFOXG1 (4 × 10^10^, 1 × 10^11^ or 2.6 × 10^11^ vg per mouse) did not significantly affect their survival compared with vehicle treatment. In the dose range finding study, there was no significant difference among the ICV treatment groups (vehicle-treated WT mice, vehicle-treated Q84P mice, and Q84P mice treated with scAAV9.hSyn1-opthFOXG1 at 4 × 10^10^, 1 × 10^11^ or 2.6 × 10^11^ vg per mouse) in probability of survival to the planned termination at 13 weeks of age. In each group of 14 to 15 animals, one to four mice were found dead prior to postnatal day 21 (P21), including one animal from the vehicle-treated WT group, three animals from the vehicle-treated Q84P group, two animals from the Q84P group treated with scAAV9.hSyn1-opthFOXG1 at 4 × 10^10^ vg per mouse, four animals from the Q84P group treated with scAAV9.hSyn1-opthFOXG1 at 1 × 10^11^ vg per mouse, and four animals from the Q84P group treated with scAAV9.hSyn1-opthFOXG1 at 2.6 × 10^11^ vg per mouse. Similarly, in a subsequent study with ICV administration of 2.6 × 10^11^ vg scAAV9.hSyn1-opthFOXG1 in Q84P animals, survival until the planned termination at 13 weeks of age was comparable across the treatment groups, with two vehicle-treated WT mice, two vehicle-treated Q84P mice and three animals from the Q84P group treated with scAAV9.hSyn1-opthFOXG1 at 2.6 × 10^11^ vg per mouse found dead before postnatal day 30 (P30).

There were no abnormal cage side observations in any group treated with scAAV9.hSyn1-opthFOXG1 (4 × 10^10^, 1 × 10^11^ or 2.6 × 10^11^ vg per mouse). Furthermore, hematoxylin and eosin staining of the brain at week 13 in the dose-range finding study revealed no scAAV9.hSyn1-opthFOXG1 treatment-related abnormalities in Q84P mice at any tested dose.

#### Body Weight

In the dose-range finding study, ICV administration of 4 × 10^10^, 1 × 10^11^ or 2.6 × 10^11^ vg per mouse of scAAV9.hSyn1-opthFOXG1 in Q84P animals and vehicle in Q84P or WT mice resulted in no significant differences in body weight among the groups (**Fig. 5A**). In a subsequent study with ICV administration of 2.6 × 10^11^ vg scAAV9.hSyn1-opthFOXG1 in Q84P mice and vehicle in Q84P or WT mice, Q84P animals treated with either vehicle or scAAV9.hSyn1-opthFOXG1 transiently weighed significantly less than WT mice treated with vehicle at 4, 5, 6 and 7 weeks of age (**Fig. 5B**). No significant differences in body weight were detected between Q84P mice treated with vehicle and those treated with scAAV9.hSyn1-opthFOXG1 at any timepoint during the study.

**Figure 5.**
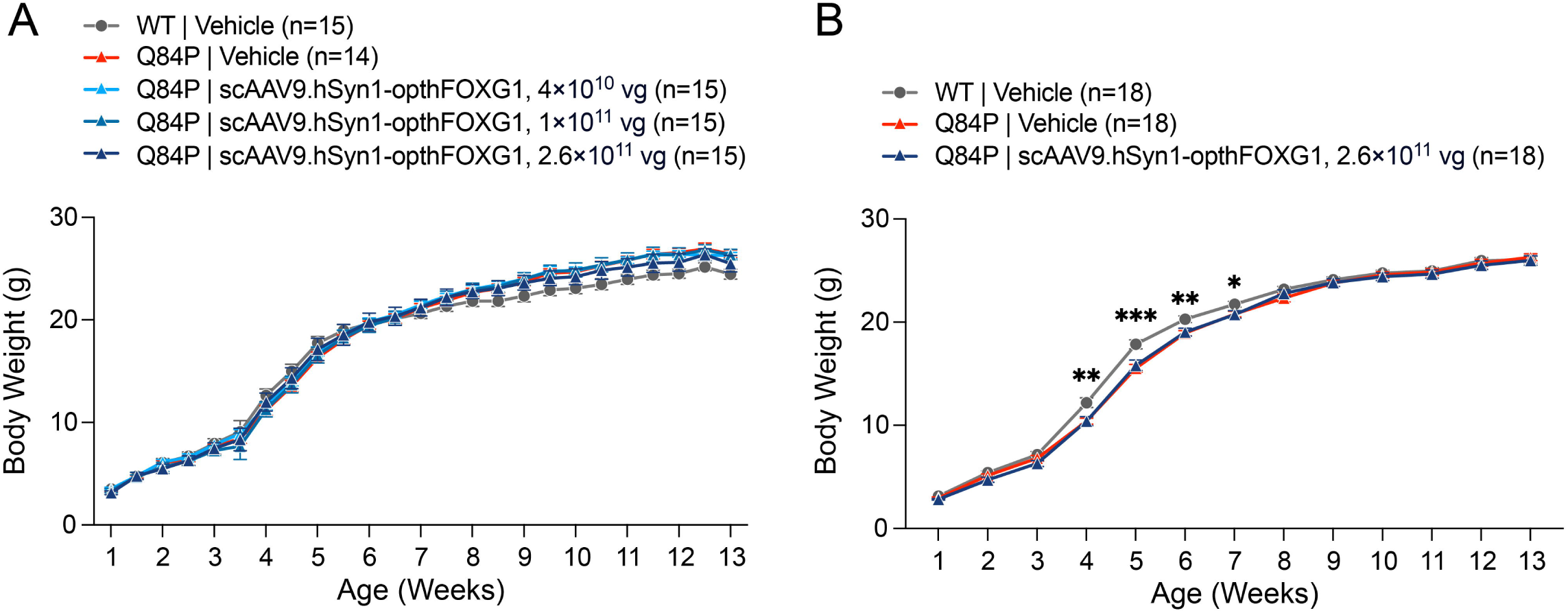
Body weights of Q84P mice treated ICV with scAAV9.hSyn1-opthFOXG1 are not significantly different from Q84P mice treated with vehicle. (**A**) In a dose-range finding study in Q84P mice, ICV administration of scAAV9.hSyn1-opthFOXG1 at 4 × 10^10^, 1 × 10^11^ or 2.6 × 10^11^ vg per mouse did not result in significant differences in body weight compared with vehicle-treated Q84P or WT mice, as assessed by two-way mixed effects ANOVA (F (4, 69) = 0.3412, p = 0.8492). Animals increased in body weight as the study progressed (F (24, 1245) = 4748, p < 0.0001). (**B**) In a study in Q84P mice with ICV administration of scAAV9.hSyn1-opthFOXG1 at a dose of 2.6 × 10^11^ vg per mouse or vehicle compared with WT mice treated with vehicle, Q84P mice treated with either vehicle or scAAV9.hSyn1-opthFOXG1 transiently weighed significantly less that WT mice treated with vehicle at 4, 5, 6 and 7 weeks of age, as assessed by two-way ANOVA with Tukey’s multiple comparison test. Means ± standard error of the mean are displayed. * p < 0.05, ** p < 0.01, *** p < 0.001

### ICV injection of scAAV9.hSyn1-opthFOXG1 in the Q84P mouse on postnatal day 6 increases FOXG1 expression in key brain regions

Vector genome (VG), human *FOXG1* mRNA and human FOXG1 protein levels confirmed delivery of scAAV9.hSyn1-opthFOXG1 to the brain of Q84P mice including cortex and hippocampus, key regions for treating FOXG1 syndrome.

In the dose-range finding study, VG levels measured at week 13 by droplet digital PCR (ddPCR) were generally dose-related in cortex and hippocampus after ICV administration of scAAV9.hSyn1-opthFOXG1, with the exception of VG levels in the cortex at the two lower dose levels. Specifically, VG levels (mean ± standard error of the mean) were 5.9 ± 2.9, 2.1 ± 0.5 and 17 ± 5.0 vector genomes/ diploid cell (VG/DC) in the cortex and 2.9 ± 0.8, 5.3 ± 1.1 and 22 ± 6.3 VG/DC in the hippocampus after ICV administration of 4 × 10^10^, 1 × 10^11^ and 2.6 × 10^11^ vg per mouse, respectively, at P6, and less than 0.05 VG/DC in all samples from vehicle-treated WT and Q84P groups. These results indicate that there was a greater than dose-proportional increase in VG levels in both cortex and hippocampus from the mid-dose to the high dose of scAAV9.hSyn1-opthFOXG1.

Human *FOXG1* mRNA levels in the Q84P mouse hippocampus and cortex, quantified by RT-qPCR and normalized to mouse beta-glucuronidase (GUSB) mRNA levels, also generally increased with dose of scAAV9.hSyn1-opthFOXG1, with the exception of human *FOXG1* mRNA levels in the cortex at the two lower dose levels. In vehicle-treated WT and Q84P mice, human *FOXG1* mRNA levels in cortex and hippocampus, normalized to mouse GUSB mRNA levels, were very low or undetectable as expected since these animals were not administered scAAV9.hSyn1-opthFOXG1. In Q84P mice treated with scAAV9.hSyn1-opthFOXG1, normalized human *FOXG1* mRNA levels in cortex and hippocampus, expressed as fold-change (mean ± standard error of the mean) relative to the average of the corresponding tissue samples from the vehicle-treated Q84P group, were 13,864 ± 6,350, 6,710 ± 2,189 and 47,217 ± 10,486 in the cortex and 45,559 ± 12,789, 89,662 ± 12,424 and 376,511 ± 112,841 in the hippocampus after ICV administration of 4 × 10^10^, 1 × 10^11^ and 2.6 × 10^11^ vg per mouse, respectively, at P6. These human *FOXG1* mRNA levels paralleled VG levels measured from the same tissue samples and suggest that there was a greater than dose-proportional increase in human *FOXG1* mRNA levels in both cortex and hippocampus from the mid-dose to the high dose of scAAV9.hSyn1-opthFOXG1.

In a subsequent study with ICV administration of scAAV9.hSyn1-opthFOXG1 at 2.6 × 10^11^ vg per mouse, VG levels (mean ± standard error of the mean) in cortex and hippocampus tissues from 11 Q84P mice that received scAAV9.hSyn1-opthFOXG1 were 7.4 ± 3.6 and 15 ± 7.4 VG/DC, respectively. In these same cortex and hippocampus samples, human *FOXG1* mRNA levels, normalized to mouse GUSB mRNA levels and expressed as fold-change relative to the vehicle treated Q84P group (mean ± standard error of the mean), were 17,585 ± 8,274 and 13,282 ± 6,089, respectively. Notably, 8 of these 11 mice had unexpectedly low levels of VG and relative human *FOXG1* mRNA in both cortex and hippocampus, less than one-tenth the levels of VG in cortex and hippocampus tissues measured in the other 3 mice from this group and less than one-tenth of the VG levels in the Q84P mouse group that received the same AAV vector and dose in the previous dose-range finding study. These results suggested that these eight Q84P mice had received lower than intended doses of scAAV9.hSyn1-opthFOXG1, so they were excluded from subsequent analyses.

Immunohistochemical staining for FOXG1 at 13 weeks of age in the dose-range finding study further confirmed delivery of scAAV9.hSyn1-opthFOXG1 to the brain in Q84P mice after ICV administration and expression of FOXG1 protein. With the low dose of scAAV9.hSyn1-opthFOXG1 (4 × 10^10^ vg per mouse), FOXG1-immunoreactive nuclei were present in neurons in the hippocampus and cerebral cortex but not in the thalamus or cerebellum. With the high dose of scAAV9.hSyn1-opthFOXG1 (2.6 × 10^11^ vg per mouse; **Fig. 6**), there was an increased number of FOXG1-immunoreactive nuclei in neurons in the telencephalon, as well as additional staining in the vestibular nuclei, inferior colliculus, and regions near the substantia nigra, compared to the vehicle-treated WT and Q84P mice.

**Figure 6.**
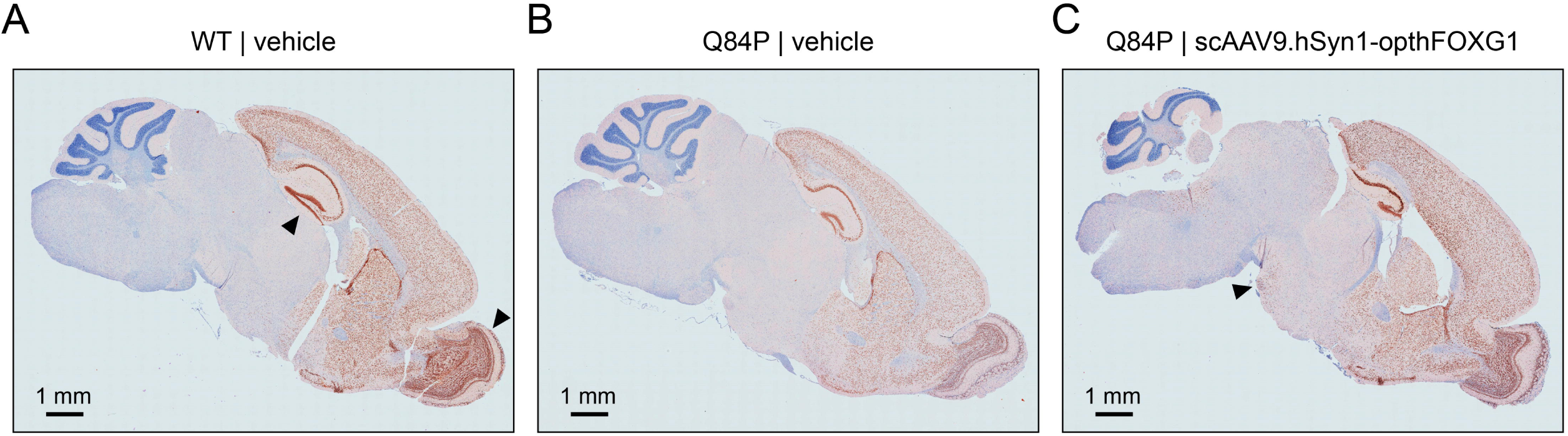
Immunohistochemical staining for FOXG1 in the brain is reduced in Q84P versus WT mice treated with vehicle and elevated after ICV treatment with scAAV9.hSyn1-opthFOXG1. Paraffin-embedded sagittal brain sections (4-5 μm thick) from mice 13 weeks of age were stained for FOXG1. (**A**) In a WT mouse (treated with vehicle), the hippocampus (arrowhead) and olfactory bulbs (arrowhead) have the highest levels of FOXG1 immunostaining in the brain. (**B**) In a Q84P mouse (treated with vehicle), the overall level of FOXG1 immunostaining in the brain is substantially reduced compared with the level in WT mice treated with vehicle. (**C**) In a Q84P mouse treated with scAAV9.hSyn1-opthFOXG1 at an ICV dose of 2.6 × 10^11^ vg, the level of FOXG1 immunostaining in the brain is higher than that in Q84P mice treated with vehicle. In particular, neuronal nuclei are labelled in the telencephalon, the vestibular nuclei, the inferior colliculus, and regions near the substantia nigra (arrowhead).

In a separate study where Q84P mice received 2.6 × 10^11^ vg scAAV9.hSyn1-opthFOXG1 by ICV injection at P6, human FOXG1 protein levels were quantified by ultra-performance liquid chromatography tandem mass spectrometry (UPLC-MS/MS) in cortex and hippocampus tissues from 13-week-old animals. In vehicle-treated Q84P and WT mice, human FOXG1 protein levels in cortex and hippocampus were below the limit of quantitation in all samples that were tested, as expected since these animals were not treated with scAAV9.hSyn1-opthFOXG1. In Q84P mice treated with scAAV9.hSyn1-opthFOXG1, human FOXG1 protein levels in cortex and hippocampus were measurable in all samples that were tested and were 0.014±0.002 and 0.016±0.004 ng/mL (mean ± standard error of the mean), respectively. The VG levels in cortex and hippocampus from these mice were 19.3 ± 4.1 and 30.6 ± 7.0 VG/DC (mean ± standard error of the mean), respectively, which overlap those in the corresponding dose group of the dose-response study.

### Some but not all behavioral deficits in Q84P mice are ameliorated by ICV treatment with AAV *FOXG1* gene replacement therapy on postnatal day 6

#### Open field activity is increased by AAV FOXG1 gene replacement therapy

Multiple open field endpoints in the dose-range finding study were significantly affected at 11 weeks of age (approximately 10 weeks post-dosing) by treatment at P6 with the top dose of scAAV9.hSyn1-opthFOXG1 (2.6 × 10^11^ vg per mouse), including total and center distance travelled and total and center rearing frequency (**Fig. 7**). There was no effect of the lower doses of scAAV9.hSyn1-opthFOXG1 (4 × 10^10^ and 1 × 10^11^ vg per mouse) on any open field parameter (**Fig. 7**). Total distance travelled at 11 weeks of age was reduced by 27% on average (not statistically significant) and increased significantly by the top dose of scAAV9.hSyn1-opthFOXG1 (2.6 × 10^11^ vg per mouse) relative to vehicle treated Q84P mice (**Fig. 7A**). Center distance travelled was significantly reduced in vehicle-treated Q84P versus WT mice at all ages evaluated (including 6 and 8 weeks of age; **SUPPLEMENTARY Fig. 8B** and **G**, respectively) and normalized at 11 weeks of age by the top dose of scAAV9.hSyn1-opthFOXG1 (2.6 × 10^11^ vg per mouse; **Fig. 7B**). An increase in average center distance travelled was also observed at 6 and 8 weeks of age with scAAV9.hSyn1-opthFOXG1 treatment of Q84P mice, though not statistically significant. Total rearing frequency at 6, 8 and 11 weeks of age showed no significant changes in vehicle-treated Q84P versus WT mice (**SUPPLEMENTARY Fig. 8C** and **8H** and **Fig. 7C**, respectively). However, total rearing frequency was significantly increased at 8 and 11 weeks of age by the top dose of scAAV9.hSyn1-opthFOXG1 (2.6 × 10^11^ vg per mouse) relative to vehicle-treated Q84P mice (**SUPPLEMENTARY Fig. 8H** and **Fig. 7C**, respectively). Center rearing frequency was reduced significantly by 77% on average at 11 weeks of age and normalized at 11 weeks of age by the top dose of scAAV9.hSyn1-opthFOXG1 (2.6 × 10^11^ vg per mouse) relative to vehicle-treated Q84P mice (**Fig. 7D**). At 6 and 8 weeks of age, center rearing frequency decreased by 48% and 55% on average, respectively, though not statistically significant (**SUPPLEMENTARY Fig. 8D** and **I**, respectively). Open field velocity showed significant reductions at 6 and 8 weeks of age (**SUPPLEMENTARY Fig. 8E** and **J**, respectively), and a 17% reduction on average at 11 weeks of age that was not statistically significant (**Fig. 7E**). Treatment with scAAV9.hSyn1-opthFOXG1 did not have a significant effect on this parameter at any dose or age tested.

**Figure 7.**
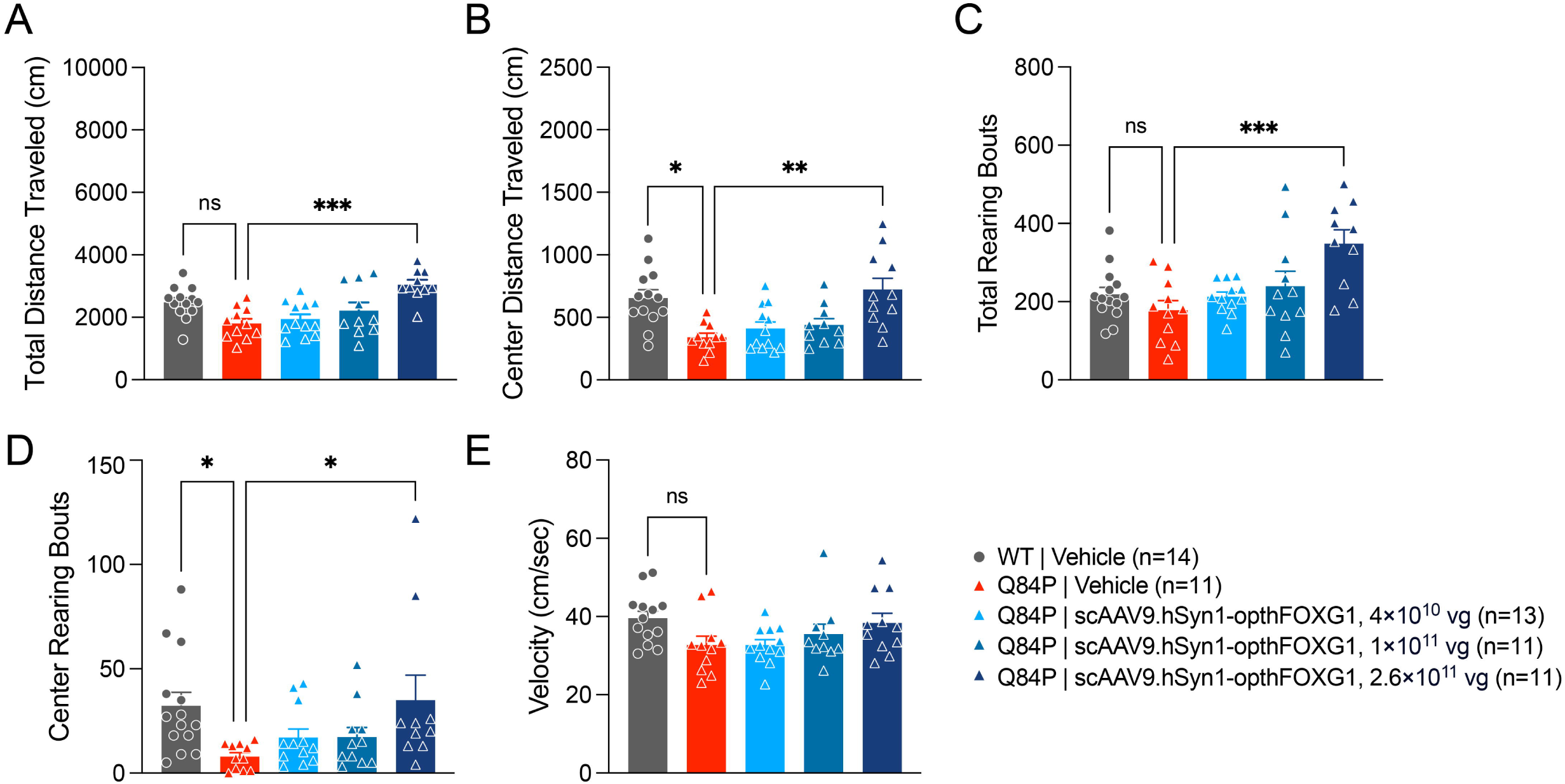
Open field center distance travelled and center rearing frequency show significant reductions in 11-week old Q84P mice compared to WT mice that are normalized by scAAV9.hSyn1-opthFOXG1 treatment at an ICV dose of 2.6 × 10^11^ vg per mouse. Decreases in other endpoints such as total distance travelled, total rearing bouts, and velocity did not reach statistical significance in vehicle-treated Q84P vs WT mice. Open field total (**A**) and center (**B**) distance travelled, total (**C**) and center (**D**) rearing frequency, and velocity (**E**) showed differences between genotypes at 11 weeks of age that were significant for center distance travelled and center rearing frequency. One-way ANOVA with Tukey’s multiple comparisons test was used to assess statistical significance. Means ± standard error of the mean are displayed. ns: not significant. * p < 0.05, ** p < 0.01, *** p < 0.001

**Figure 8.**
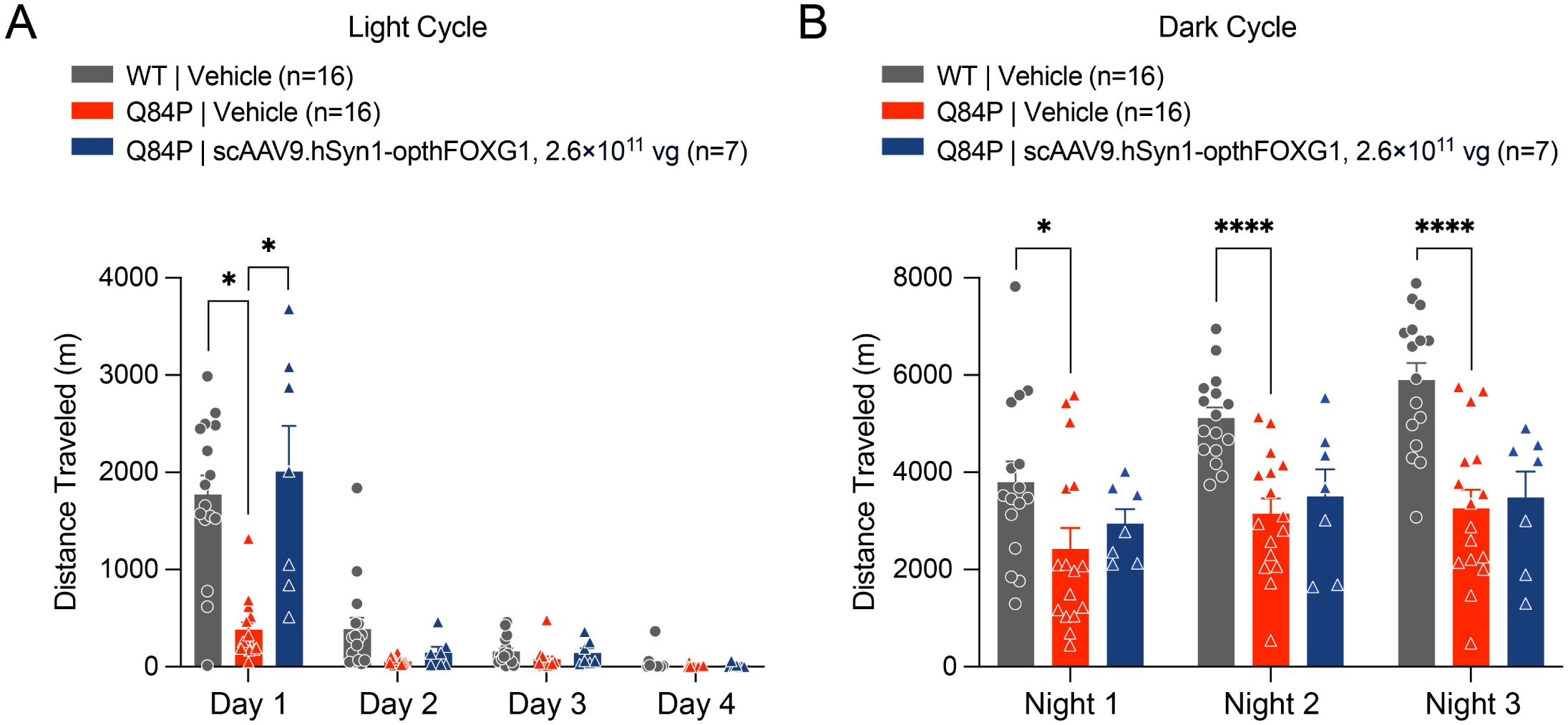
Running wheel total distance travelled during the light and dark cycles at 12 weeks of age, excluding Q84P mice treated with scAAV9.hSyn1-opthFOXG1 that had less than one-tenth of expected vector genome levels in cortex and hippocampus tissues. The total distances travelled during the animals’ light (from 7AM to 7PM) and dark (from 7PM to 7AM) cycles, assessed on 4 days and 3 nights, respectively, showed significant differences between the treatment groups, as assessed by two-way ANOVA (Genotype X Time Interaction: F (37, 149) = 5.36, p < 0.001). (**A**) The total distance travelled during the animals’ light cycle (from 7AM to 7PM) show a significant reduction in Q84P mice compared with WT mice treated with vehicle on Day 1 of assessment, with a significant normalization by treatment with scAAV9.hSyn1-opthFOXG1, as assessed with Tukey’s multiple comparison post-hoc test. (**B**) The total distance travelled during the animals’ dark cycle (from 7PM to 7AM) showed significant reductions in Q84P mice compared with WT mice treated with vehicle on Night 1, Night 2 and Night 3 of assessment, as assessed with Tukey’s multiple comparisons post-hoc test. There was no significant effect on running wheel distances travelled on any night of assessment, of treatment with scAAV9.hSyn1-opthFOXG1. Means ± standard error of the mean are displayed. * p < 0.05, **** p< 0.0001

In the single dose study with scAAV9.hSyn1-opthFOXG1 (2.6 × 10^11^ vg per mouse), Q84P mice at 11 weeks of age treated with vehicle travelled significantly less center distance (**SUPPLEMENTARY Fig. 9B** and **F**) and reared significantly more overall (**SUPPLEMENTARY Fig. 9C** and **G**) compared with WT mice treated with vehicle. When all Q84P mice treated with scAAV9.hSyn1-opthFOXG1 were included in the analysis (**SUPPLEMENTARY Fig. 9B** and **C**), there were no significant treatment effects compared with vehicle-treated Q84P mice in either parameter. However, when the eight Q84P mice that had less than one-tenth of expected vector genome levels in cortex and hippocampus tissues were excluded from the analysis (**SUPPLEMENTARY Fig. 9F** and **G**), there was a significant increase in center distance travelled and total rearing frequency after scAAV9.hSyn1-opthFOXG1 versus vehicle treatment in Q84P mice. In this study, total distance travelled (**SUPPLEMENTARY Fig. 9A** and **E**) and center rearing frequency (**SUPPLEMENTARY Fig. 9D** and **H**) were not significantly different between Q84P and WT mice treated with vehicle. However, scAAV9.hSyn1-opthFOXG1-treated Q84P mice exhibited significantly greater total distance travelled than those treated with vehicle, when the eight Q84P mice that had less than one-tenth of expected VG levels in cortex and hippocampus were excluded (**SUPPLEMENTARY Fig. 9E**).

Our findings generally demonstrate beneficial effects on open field activity of the top ICV dose of scAAV9.hSyn1-opthFOXG1 (2.6 × 10^11^ vg per mouse) at 11 weeks of age, but generally not at 6 or 8 weeks of age, or at 4 × 10^10^ or 1 × 10^11^ vg per mouse dose levels at any age tested. These results suggest that the effects of scAAV9.hSyn1-opthFOXG1 administration require events downstream of exogenous FOXG1 expression that occur over 10 but not 7 or fewer weeks following treatment.

#### Running wheel activity on Day 1 of assessment is increased by AAV FOXG1 gene replacement therapy

Running wheel activity was evaluated at 12 weeks of age during the animals’ light cycle (from 7AM-7PM) and dark cycle (from 7PM-7AM) in the single dose study with 2.6 × 10^11^ vg scAAV9.hSyn1-opthFOXG1. During the day cycle, Q84P mice treated with vehicle travelled significantly less than WT mice treated with vehicle on the first day of assessment whether all mice were analyzed (**SUPPLEMENTARY Fig. 10A**) or the eight Q84P mice with less than one-tenth of expected VG levels were excluded (**Fig. 8A**) and on the second day of assessment when all mice were analyzed (**SUPPLEMENTARY Fig. 10A**), but not on day 3 or 4 with either analysis (**Fig. 8A** and **SUPPLEMENTARY Fig. 10A**) . In both analyses, there was a significant restoration of running wheel activity on day 1 of assessment after ICV treatment of Q84P mice with scAAV9.hSyn1-opthFOXG1 compared with vehicle. There was no significant effect of scAAV9.hSyn1-opthFOXG1 treatment on the day 2 running wheel deficit in all mice (**SUPPLEMENTARY Fig. 10A**). Notably, day 1 is a time period in the running wheel test that is associated with acclimating to and exploring a novel environment (Santos *et al.* 2022), suggesting that AAV *FOXG1* gene replacement therapy improves habituation to and exploring a new environment.

Q84P mice treated with vehicle also travelled significantly less than WT mice treated with vehicle during the dark cycle on the first (excluding eight Q84P mice with less than one-tenth of expected VG levels), second and third nights of testing (**Fig. 8B** and **SUPPLEMENTARY Fig. 10B**). Whether all Q84P mice treated with scAAV9.hSyn1-opthFOXG1 were included in the analysis (**SUPPLEMENTARY Fig. 10B**) or the eight Q84P mice that had less than one-tenth of expected VG levels in cortex and hippocampus were excluded from the analysis (**Fig. 8B**), there was no treatment effect detected on any of the nights tested.

In summary, the Q84P mice travelled significantly less on the running wheel than WT mice treated with vehicle on Day 1 of assessment and Nights 2 and 3 of assessment. There was a beneficial effect of AAV *FOXG1* gene replacement therapy with scAAV9.hSyn1-opthFOXG1 in Q84P mice on Day 1 of assessment only, with a normalization of running wheel distance travelled.

#### Several SmartCube® behavioral endpoints are improved by AAV FOXG1 gene replacement therapy

Phenotypic profiling in SmartCube® provides an assessment of approximately 2,000 features per animal per timepoint, where the features include spontaneous behaviors and responses to challenges. Over the course of the 45-minute testing session, a total of more than 100,000 features are evaluated. At 13 weeks of age, there were significant differences between Q84P and WT mice treated with vehicle (**Fig. 9A**). Small but significant recovery was detected in Q84P mice treated ICV with 2.6 × 10^11^ vg scAAV9.hSyn1-opthFOXG1 compared with Q84P mice treated with vehicle (**Fig. 9A**).

**Figure 9.**
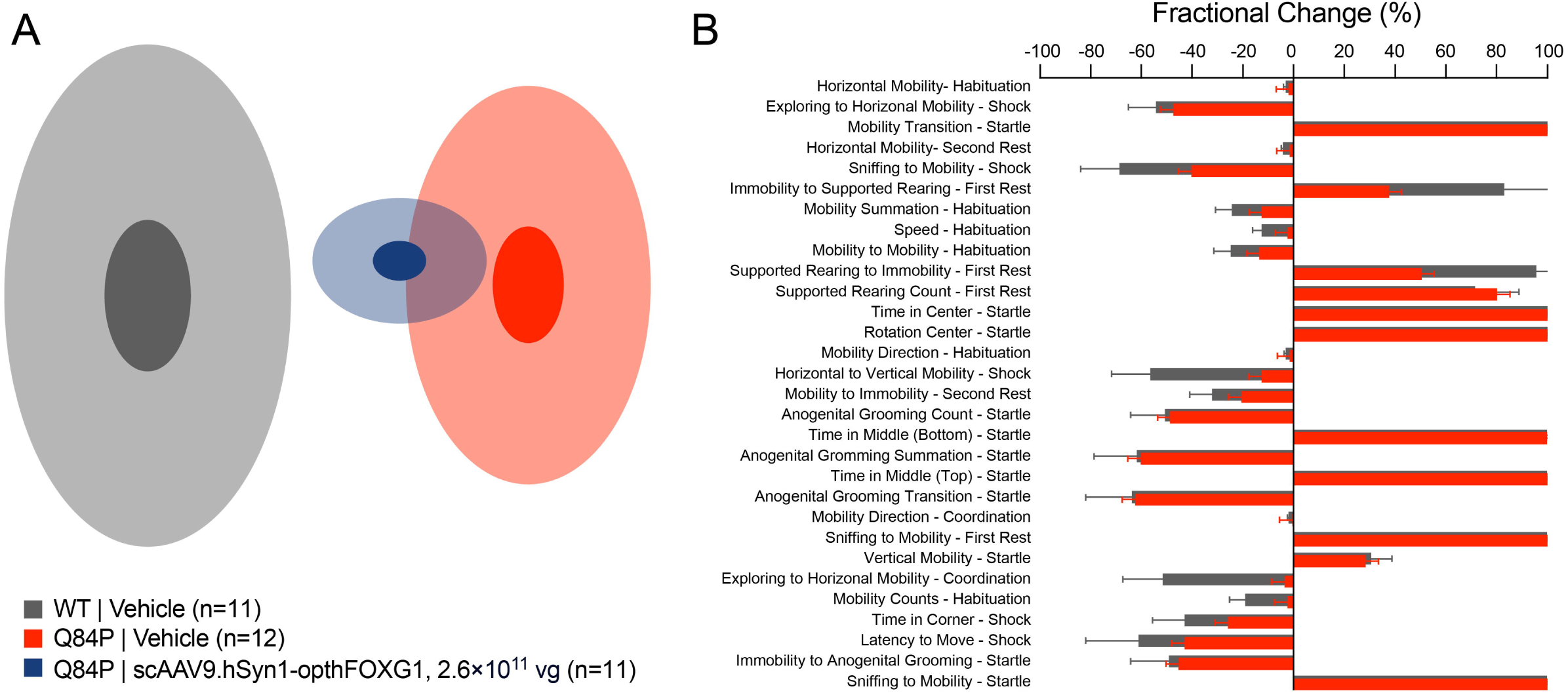
SmartCube^®^ spontaneous behaviors and responses to challenges at 13 weeks of age. (**A**) DRFA (de-correlated ranked feature analysis) cloud analysis. There were significant differences between Q84P and WT mice treated with vehicle (Discrimination = 97.1%, p < 0.0001) and significant recovery in Q84P mice treated ICV with 2.6 × 10^11^ vg scAAV9.hSyn1-opthFOXG1 versus vehicle (Recovery = 23.2%, p = 0.0160). (**B**) Thirty top features from the SmartCube^®^ results shown as fractional (%) change versus the WT vehicle group. Larger absolute fractional changes indicate greater differences from the WT vehicle group whereas smaller absolute fractional changes indicate smaller differences from the WT vehicle group. Four animals per group were reserved for other analyses and not tested in SmartCube^®^ and SmartCube^®^ data were lost from one vehicle-treated WT mouse. Means ± standard error of the mean are displayed.

Of the 30 top features of the SmartCube® that were significantly different between Q84P and WT mice treated with vehicle, six features were robustly improved by ICV treatment with 2.6 × 10^11^ vg scAAV9.hSyn1-opthFOXG1: Sniffing to Mobility transition during the Shock phase, Immobility to Supported Rearing transition during the first rest, Supported Rearing to Immobility transition during the first rest, Horizontal to Vertical Mobility transition during the Shock phase, Exploring to Horizontal Mobility transition during the Coordination task, and Mobility Counts in the Habituation phase (**Fig. 9B**). In general, the features that were improved by AAV *FOXG1* gene replacement therapy with scAAV9.hSyn1-opthFOXG1 reflect an improvement in habituation to a novel environment.

#### Contextual fear conditioning is not affected by AAV FOXG1 gene replacement therapy

The contextual fear conditioning phenotype in 12-week-old Q84P mice, comprising a decrease in freezing, was not affected by scAAV9.hSyn1-opthFOXG1 administration at any ICV dose tested in the dose-range finding study (**Fig. 10**).

**Figure 10.**
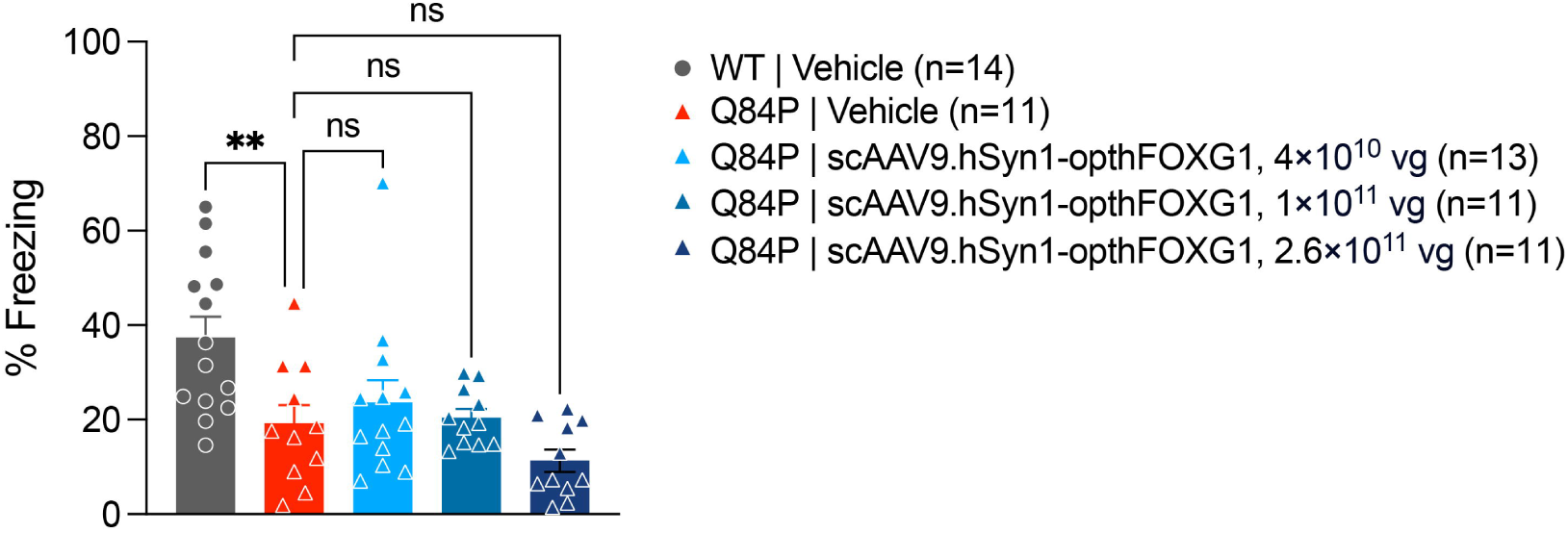
Contextual fear conditioning showed a significant difference between vehicle-treated Q84P and WT mice at 12 weeks of age, but no effect of ICV treatment of Q84P mice with scAAV9.hSyn1-opthFOXG1 at any dose tested (4 × 10^10^, 1 × 10^11^ or 2.6 × 10^11^ vg per mouse), as assessed by one-way ANOVA with Tukey’s multiple comparisons post-hoc test. Means ± standard error of the mean are displayed. ns: not significant. ** p < 0.01

In the dose-range finding study and/or the single dose study, there were no phenotypic differences between Q84P and WT mice treated with vehicle in cued fear conditioning (dose-range finding study), grip strength (dose-range finding study), wire hang (single dose study) or tapered balance beam (single dose study), precluding the evaluation of an effect of scAAV9.hSyn1-opthFOXG1 administration on phenotypic differences in these behaviors (data not shown).

## DISCUSSION

To demonstrate the therapeutic potential of AAV gene therapy to treat FOXG1 syndrome, it is critical to establish safety and functional efficacy in an animal model of disease. Here, in a new patient-specific mouse model of FOXG1 syndrome that expresses Q84P mutant FOXG1, we showed that AAV *FOXG1* gene replacement therapy to deliver human FOXG1 to neurons is well-tolerated and ameliorates multiple behavioral deficits that collectively reflect improvements in anxiety and habituation to a novel environment. Specifically, ICV administration of scAAV9.hSyn1-opthFOXG1 at P6 in heterozygous Q84P mice has beneficial effects on a number of open field, running wheel (only day 1 of assessment) and SmartCube^®^ phenotypes. However, there was no effect on the contextual fear conditioning deficit. All tested ICV doses of scAAV9.hSyn1-opthFOXG1, including the efficacious top dose, were well-tolerated with no significant effects on body weight, no abnormal cage side observations and no microscopic abnormalities in brain structure. Vector genome, human *FOXG1* mRNA and human FOXG1 protein levels in cortex and hippocampus, two key brain regions relevant to FOXG1 syndrome, were measurable and generally dose-related after ICV administration of scAAV9.hSyn1-opthFOXG1.

Our new Q84P mouse model of FOXG1 syndrome shares several behavioral features with another Q84P mouse model of FOXG1 syndrome reported recently by Jeon *et al.* (2025). In particular, movement disorders, repetitive behaviors and anxiety were observed by Jeon *et al.* (2025) based on reduced wire hang time, increased repetitive grooming behavior, and decreased open field activity, respectively. In our model, a consistent reduction in open field activity was also observed, as well as a consistent decrease in running wheel distance on only the day 1 assessment and changes in SmartCube® endpoints that collectively are consistent with an increase in anxiety and a decrease in habituation to a novel environment. However, we did not observe a change in wire hang time in our Q84P mouse model, in contrast to Jeon *et al.* (2025). The behavioral deficit of decreased open field activity that is present in both Q84P mouse models reinforces this phenotype as a consequence of replacing one *Foxg1* allele with mutant Q84P *Foxg1*. Furthermore, the increase in anxiety and decrease in habituation to a novel environment that are reflected by reduced open field activity, correspond to key clinical manifestations of FOXG1 syndrome. Therefore, both Q84P mouse models recapitulate important functional aspects of FOXG1 syndrome and provide useful models of disease to evaluate effects of potential therapeutics on these behavioral deficits.

In the only publication to date on the *in vivo* efficacy of AAV gene therapy in FOXG1 syndrome, a mouse model of FOXG1 haploinsufficiency that does not express mutant FOXG1 (*Foxg1fl/+;NexCre* mouse, Jeon *et al.* 2024) was used to evaluate neuroanatomical effects in the brain. ICV administration of AAV9.CBA.hFOXG1 on postnatal day 1 corrected or improved some neuroanatomical deficits in the brain as assessed on postnatal day 27, increasing the reduced numbers of neurons in the cortex and oligodendrocyte progenitor cells in the corpus callosum, increasing the reduced myelination in the corpus callosum, and improving hippocampus dentate gyrus structural abnormalities. However, there was no change in the decreased thickness of the cortex. The improvements in neuroanatomical deficits, while encouraging, do not address the effect of AAV *FOXG1* gene replacement therapy on functional endpoints. Moreover, the lack of mutant FOXG1 in the *Foxg1fl/+;NexCre* mouse may impact the effect of a potential therapeutic approach. For example, if there is a dominant negative effect of mutant FOXG1, then the *Foxg1fl/+;NexCre* mouse would not enable an appropriate evaluation of the effectiveness of a potential therapeutic approach. Our studies of AAV *FOXG1* gene replacement therapy, with ICV administration of scAAV9.hSyn1-opthFOXG1 in a patient-specific Q84P mouse model that expresses mutant FOXG1, demonstrate improvements in some but not all behavioral deficits, and in particular show that poor habituation to a novel environment and increased anxiety are ameliorated.

Elevated levels of FOXG1 expression are associated with an autistic spectrum disorder similar to FOXG1 syndrome (Yeung *et al.* 2009, Brunettie-Pierri *et al.* 2011). Therefore, it is particularly important to ensure that there are no detrimental effects of AAV *FOXG1* gene replacement therapy, which may result in long-term levels of exogenous human FOXG1 that are significantly higher than normal. In our studies with 12 weeks of observation in heterozygous Q84P mice following ICV administration of scAAV9.hSyn1-opthFOXG1, there were no gene therapy-related adverse effects on survival or body weight, cage side observations, neuropathology in the brain, or behavioral phenotype. Supraphysiological FOXG1 levels from AAV or other therapeutic approaches may be beneficial and differentially impactful depending on treatment timing with respect to developmental stage and targeted cell type for expression. However, the potential risks associated with continuous elevated levels of FOXG1 expression need to be explored further with comprehensive toxicity studies together with quantitation of levels of exogenous FOXG1 protein in the brain for correlation with safety and any adverse findings.

In the other Q84P mouse model of FOXG1 syndrome (Jeon *et al.* 2025), the mutant Q84P FOXG1 protein was reported to interact with full-length FOXG1 and sequester it into distinct subcellular domains. A dominant-negative effect of mutant Q84P FOXG1 protein would have important implications for therapeutic approaches to treating FOXG1 syndrome, in that elevating levels of WT FOXG1 protein may or may not be sufficient to overcome the effects of the mutant protein. Our studies in a Q84P mouse model of FOXG1, demonstrating efficacy of AAV *FOXG1* gene replacement therapy on several behaviors related to anxiety and habituation to a novel environment, suggest that for these behaviors, increasing levels of WT FOXG1 are sufficient to overcome the effects of haploinsufficiency and/or expression of mutant Q84P FOXG1 for potential therapeutic benefit. However, contextual fear conditioning deficits were not altered by AAV *FOXG1* gene replacement therapy with ICV administration of scAAV9.hSyn1-opthFOXG1. It remains to be determined if this and other behavioral deficits can be addressed by replacement of missing FOXG1 protein earlier in development or in other cell types, or may require suppression of mutant Q84P FOXG1 protein.

In summary, our experiments with ICV administration of scAAV9.hSyn1-opthFOXG1, in a patient-specific mouse model of FOXG1 syndrome that contains a mutant Q84P *Foxg1* allele in place of a WT *Foxg1* allele, demonstrate the potential therapeutic benefit of AAV *FOXG1* gene replacement therapy in neurons for the treatment of FOXG1 syndrome. In particular, ICV administration of scAAV9.hSyn1-opthFOXG1 was well-tolerated and several behaviors related to anxiety and habituation to a novel environment were improved, suggesting that for these behaviors, increasing levels of WT FOXG1 are sufficient to overcome the effects of haploinsufficiency and/or expression of mutant Q84P FOXG1. Further studies in the Q84P mouse model of FOXG1 syndrome to explore efficacy of AAV *FOXG1* gene replacement therapy when administered at other ages or to other cell types, or in conjunction with inhibitors of mutant Q84P FOXG1 protein, may be useful for evaluating additional benefits of this therapeutic approach.

## MATERIALS AND METHODS

### Constructs and vectors

scAAV9.hSyn1-opthFOXG1 was produced by the University of Massachusetts Chan Medical School Viral Vector Core. The transgene DNA was packaged in a recombinant AAV9 capsid. Recombinant AAV vectors were produced by triple transfection of adherent HEK293 cells and purified using previously described protocols. In brief, recombinant vectors were purified from clarified lysates using cesium chloride density gradient centrifugation. Vectors were formulated in phosphate-buffered saline (PBS) with 5% sorbitol and 0.001% (w/v) Pluronic F-68^®^, pH 7.2-7.4, and stored below -60°C. Vector titers were measured by ddPCR. For vectors used for in vivo studies, the relative purity of the capsid proteins was confirmed using sodium dodecyl-sulfate polyacrylamide gel electrophoresis (SDS-PAGE) and protein staining. On the day of use, AAV vectors were removed from storage at -60°C, allowed to thaw on refrigerated cold packs, and then maintained at approximately 4°C. Dilutions were made with formulation buffer and then stored at approximately 4°C for no more than 1 week until use. Vehicle groups received formulation buffer.

### Animals and treatments

#### Animals

All animal procedures were approved by the Institutional Animal Care and Use Committee (IACUC).

Q84P heterozygous mice were generated by Dr. Teresa Gunn using CRISPR/Cas9 gene editing. A ribonucleoprotein mix consisting of two sgRNAs (sequences: CCCGGCCCCGCAGCCCCCGC and GGGGCGCCGCGCGCCUGCGG, Synthego Corp.) and SpCas9 2NLS nuclease (Synthego Corp.) was combined with a single stranded Alt-R HDR Donor Oligo from Integrated DNA Technologies (sequence: CAGCAGCAGCAGCAGCCGCCCCCGGCCCCGCAGCCCCCGCCAGGCGCGCGGCGCCCCAGCAGCCGACGACGACAAGGGTCC) and electroporated into 1-cell C57BL/6J embryos. All mice used in the studies described here were descended from a single male founder and had been backcrossed to wildtype C57BL/6J for at least 5 generations.

Male heterozygous Q84P mice were received at PsychoGenics from Dr. Teresa Gunn and bred with WT C57BL/6 female mice. At birth, a tail snip sample was taken for genotyping to identify heterozygous or WT animals to be placed on study. All animals were examined, manipulated and weighed prior to initiation of the study at postnatal day 6 to assure adequate health and suitability and to minimize non-specific stress associated with manipulation. Mice were maintained on a 12-hour light/dark cycle with food and water available ad libitum.

During the study, animal body weights were recorded daily or twice per week until weaning and twice per week or weekly after weaning, and survival was checked twice per day.

#### Genotyping

Genomic DNA was isolated from neonate tail clippings using Zymo Quick DNA 96 Plus Kit (Cat# D4070) according to the manufacturer’s instructions. PCR reactions were set up using QuantaBio’s sparQ HiFi PCR Master Mix (Cat# 95192-050) in 25µL reaction volume, with the following primers: Foxg1_Q86_253F (5’-CCA AGT CCT CGT TCA GCA TC3’) and Foxg1_Q86_856R (5’TGC TTG TTC TCG CGG TAG TA-3’). The PCR parameters were 94°C 3 min; [94 °C 30s, 60 °C 35s 72 °C 35s] x35 cycles; 72°C 5 min; 12 °C hold. A sample of the PCR reaction was then used for electrophoresis to ensure that a dominant approximately 600bp DNA amplicon was present in each loaded lane. Remaining PCR reactions were cleaned up using ThermoFisher’s Exosap-IT PCR Product Cleanup reagent (Cat# 78201.1.ML) and then a sample was prepared with sequencing primer (Foxg1_Q86_856R (5’TGC TTG TTC TCG CGG TAG TA3’) for shipment to GeneWiz for DNA sequencing. DNA sequence files were then reviewed individually to assign genotype.

#### Test articles and dosing

Stock solutions of scAAV9.hSyn1-opthFOXG1 (2.69 × 10^13^ GC/mL) and formulation buffer (vehicle: PBS with 5% sorbitol and 0.001% F-68) were stored at -80°C. On the first day of dose administration, test articles were thawed on wet ice. Once thawed, test articles were stored at 4°C and did not undergo any additional freeze-thaw cycles.

For administration of either vehicle or scAAV9.hSyn1-opthFOXG1, animals were anesthetized using cryoanesthesia. ICV injections were administered bilaterally (5 µL/side for P6 mice) with 10 µl Hamilton Syringes (1701 RN Model, Cat# 7653-01) and custom Hamilton 30 ½ g needles (point style 4, bevel 12°, Cat# 7803-07). Animals were then placed on a warm heating pad immediately following the ICV injection. Once all animals from the litter were dosed and warmed, they were returned to the dam.

### Behavioral assessments

#### Neonatal well-being indices

The following well-being indices were evaluated every 48 hours from birth through P16, with the exception of milk content, which was assessed through P6, and respiration rate, which was assessed every four days until P20: gasping (yes/no scoring; presence or absence of mouth and thoracic movements indicative of gasping), skin color (0-2 scoring; 2 indicates a pink, well-oxygenated color while 0 indicates a pale bluish cast), milk content (0-2 scoring; 2 indicates normal milk content), muscle tone (0-2 scoring; 2 indicates a strong general body tonus when the animal was lifted and strong resistance of a hindlimb when the limb was flexed, whereas 0 indicates a limp or flaccid tonus), and respiration rate (1-3 scoring; 2 indicates normal respiration, 3 indicates elevated respiration, and 1 indicates depressed respiration). Body temperature was also taken after respiration assessment.

#### Neonatal motor tests

*Righting reflex*. The righting reflex assessment was performed every 48 hours through P14. Animals were placed on their backs and were given a total of 30 seconds to right themselves. The latency to right was recorded.

*Geotaxis*. The geotaxis assessment was performed every 48 hours through P14 to measure motor coordination and function of the vestibular system through the ability of the animal to orient itself when placed face down on an inclined platform. Animals were given a maximum of 60 seconds to complete the test. The latency to turn 180° was recorded.

*Tube test*. The tube test for hindlimb strength and general body muscle tone assessment was performed every 48 hours through P14. The test was performed in 2 consecutive trials. In each trial, the mouse was placed face down, hanging by its hind-limbs in a plastic 50ml centrifuge tube. Latency to fall into the tube was recorded.

#### Ultrasonic vocalization

The USV assessment was performed at P9. Briefly, the cage with the dam and the litter was taken to the testing room. The dams were taken out of the cages at least 30 min prior to test. Pups remained in the home cage with familiar bedding and nest material, and then were placed inside a warmed cage (34°C) for a 20 min adaptation period. During the test, one pup at a time was taken to a clean Plexiglas chamber for a 5 min observation at a temperature of 30°C. The isolation response was measured with an ultrasound microphone placed approximately 6 cm above the pup. The USVs were recorded through microphones and captured by NOLDUS software.

#### Open field

The open field test was performed at 4, 6, 8 and 11 weeks of age, using plexiglass square chambers (27.3 x 27.3 x 20.3 cm; Med Associates Incs., St Albans, VT) surrounded by infrared photobeam sources (16 x 16 x 16). The enclosure was configured to split the open field into a center and periphery zone and the photocell beams were set to measure activity in the center and in the periphery of the chambers. Horizontal activity (distance traveled) and vertical activity (rearing) were measured from consecutive beam breaks. Total ambulatory distance, ambulatory distance in center, total rearing, and rears in the center were measured over 30 minutes.

#### Clasping

Clasping was assessed at 4, 6, 8, 10, 11, and 12 weeks of age. The animal was suspended by the tail for 30 seconds and clasping behavior was scored. A score of 0 indicated the animal had a normal splay, with no clasping present. A score of 1 indicated that the animal displayed a forelimb and/or hind limb clasping, with limbs pulled tightly to the core.

#### Grip strength

Grip strength was evaluated at 4, 6, 8, 10, 11, and 12 weeks of age. Mice were held by the tail and lowered toward the mesh grip piece on the push-pull gauge (San Diego Instruments, San Diego, CA) until the animal grabbed with all four paws. The animal was lowered toward the platform and gently pulled backwards with consistent force by the experimenter until the animal released its grip. The combined grip force of all four limbs was recorded on the strain gauge. Animals were assessed across five consecutive trials.

#### Wire hang

The wire hang test was conducted at 10 weeks of age, using a modified protocol as described in Santa-Maria *et al.* (2012). Mice were placed on top of a standard wire cage lid. The lid was lightly shaken to cause the animal to tighten its grip and the lid was then turned upside down. The latency of mice to fall off the wire grid was measured, and average values were computed from three trials (30 seconds apart). Trials were stopped if the mouse remained on the lid after 5 minutes.

#### Tapered balance beam

The tapered balance beam assessment was performed at 11 weeks of age. The balance beam consisted of a strip of smooth black acrylic 100 cm in length, with a square cross section that tapers from a width of 1.5 cm to 0.5 cm. The beam also consists of a 0.5 cm safety ledge located 2 cm below the beam. The ledge maintains a constant width of 0.5 cm as the beam tapers. The angle of the beam is 17° from horizontal running from low to high. The highest point of the beam is approximately 58 cm from the floor. At the opposite side of the balance beam (‘end’ portion) there is a goal box which rests on the aforementioned support stand. The goal box is constructed from black acrylic, measuring 10.5 cm^3^ and containing a 3 cm² entrance hole.

On the training day, each animal completed 4 traversals in order to be considered trained for the beam. On testing day (24 hours later), mice received 3 trials of testing with an inter trial interval (ITI) of 30 seconds. Mice were placed on the bottom of the beam, facing away from the goal box. The time from placement on the beam to turning to face the goal box was recorded as the latency to turn. The maximum amount of time an animal had to complete the turn was 120 seconds. If the animal was unable or refused to turn after 120 seconds, it was positioned on the beam facing the goal box for the next phase of the experiment. Once the animal was facing the goal box, the latency to traverse the beam was recorded. The maximum amount of time an animal had to complete the traversal was 120 seconds. During the beam traversal, the number of foot slips was recorded. The slip-step ratio was calculated by dividing the total number of slips by the total number of steps taken per limb.

#### Running wheel

Running wheel activity was assessed at 12 weeks of age. The wheels used were from Lafayette Instrument®, Mouse Activity Wheel with Dual Licometer (model 80822). The polycarbonate chambers measured 13.9”L x 9.25”W x 7.875”H (35.3 23.5 x 20 cm) and the aluminum running wheels measured 5.0” ID (12.7 cm) by 2.25” (5.72 cm) width (inside) for a Run Distance of 0.40 meters/revolution. The run surface consisted of 38 rods 0.188” diameter on 0.4298” centers with a 0.2418” gap. The metric equivalent is approximately 4.8 mm diameter on 10.9 mm centers with a 6.14 mm gap. The activity wheels were connected to an interface which electronically recorded the animal’s activity (running interval, average speed, total distance).

Animals were loaded into the chambers containing the running wheels at approximately 11AM on day 1 of assessment and subsequently removed from the chambers at the same time on day 4, 72 hours later. Mice remained in the activity wheel chambers 24 hours/day the three consecutive testing days, after which they were returned to their home cages.

#### Acoustic startle

The acoustic startle test was administered at 6 and 8 weeks of age. Mice were placed in the startle enclosures and secured in the sound-attenuated startle chamber (Med Associates Inc., St Albans, VT) on top of a force transducer plate that measures the force of the movements made by the mouse for a 5 min habituation period of white noise (70 dB). Subsequent test sessions consisted of 10 blocks of eleven trials each. Within each block, stimuli of 70, 75, 80, 85, 90, 95, 100, 105, 110, 115, and 120 dB were presented in a random order with a variable inter-trial interval of 15 sec on average (range: 10-20 sec). The duration of the stimulus was 40 milliseconds (ms). Responses were recorded for 150 ms from startle onset and were sampled every ms.

#### Fear conditioning

The fear conditioning test was conducted at 10 weeks of age using the fear conditioning system manufactured by Coulbourn Instruments (PA, USA).

On day one 1, mice were placed into the conditioning chambers to habituate to the context for 120 sec where they were exposed to a three 20 second 80 dB tone (conditioned stimulus, CS) spaced 100 seconds apart. Fifteen seconds after each tone, mice received a foot shock (0.5 mA for 1 sec), the unconditioned stimulus (US). The mouse remained in the conditioning chamber for another 60 sec and then returned to its home cage. Twenty-four hours after training, the mice were tested for contextual memory where they were placed into the same chamber they were trained in for a period of 5 min without shock or any other interference.

Twenty-four hours after contextual fear conditioning, animals were tested for cued memory. Mice were placed in a novel context for 2 min (Pre-Cue). Then the CS (80dB tone) was presented for 3 minutes. Freezing behavior, defined as the complete lack of movement, was captured automatically with a video system and FreezeView software (Coulbourn Instruments, PA, USA).

#### Y-Maze

The Y-maze was performed at 9 weeks of age. Spontaneous alternation was evaluated in a single 10 min trial in a Y-maze consisting of three identical arms (40 x 10 x 20 cm) and each mouse was allowed to see distal spatial landmarks. The animal was placed in the middle of the three arms and allowed to explore freely. Arm entry was considered successful when hind paws were placed in the arm in full. Spontaneous alternation was described as successive entries into three arms, in overlapping triplet sets. The effect was calculated as percent alternation = [no. of alternations / (total number of arm entries± 2)] x 100. Total entries were recorded as an indication of ambulatory activity and mice that performed fewer than 12 entries in 10 min were excluded from the analysis.

#### SmartCube^®^

The SmartCube^®^ assessment was performed at 13 weeks of age. SmartCube^®^ is a high through-put proprietary platform developed by PsychoGenics that employs computer vision to detect changes in body geometry, posture, and behavior (both spontaneous and in response to specific challenges).

Mice were taken in their home cage to the SmartCube^®^ suite of experimental rooms where they remained until they were placed in the apparatus. The standard SmartCube^®^ protocol was run for a single session lasting 45 minutes. After the session, mice were placed back into their home cage and were returned to the colony room.

### Plasma, serum, cerebrospinal fluid and tissue collection

For plasma collection at 7 weeks of age, submandibular blood collection was performed.

For plasma and serum collection at 12 weeks of age, cardiac puncture was performed. Blood was processed to plasma using K-EDTA tubes or to serum using serum collector tubes, and then stored at -80°C.

For CSF collection, animals at 12 weeks of age were anesthetized with isoflurane and placed in a stereotaxic frame for collection of approximately 2µl of CSF by cisternae magna puncture.

After blood and/or CSF collection, animals were perfused with ambient temperature PBS to remove blood cells. The brain was then removed and either dissected to collect the left and right cortices and hippocampi to snap-freeze, or hemisected to drop fix the left hemisphere and collect and snap-freeze the right cortex and hippocampus. After snap-freezing, tissue samples were stored at -80°C. For drop fixation, the left brain hemisphere was transferred to freshly prepared 4% phosphate buffered paraformaldehyde overnight at 4°C, and then changed to PBS and stored at 4°C.

### Vector genome and human FOXG1 mRNA quantitation

#### DNA and RNA isolation

DNA and RNA samples were prepared with the Qiagen AllPrep DNA/RNA Kit for 96-well format. Frozen tissue from mouse cortex (approximately 75 mg) or hippocampus (approximately 15 mg) was homogenized in 700 mL Buffer RLT + 10 µL per mL β-mercaptoethanol using the FastPrep-24 (6m/s for 20 seconds per cycle for a total of 2 cycles), then frozen for storage. Immediately before processing, tissue homogenate samples were thawed at room temperature and centrifuged. Supernatants were then processed according to the Qiagen AllPrep DNA/RNA isolation kit’s protocol (Simultaneous Purification of DNA and RNA from Tissues using Spin Technology). In order to process less than 10mg of tissue per Qiagen’s recommendations, cortex supernatant was diluted 1:5 in Buffer RLT+β-mercaptoethanol (hippocampus supernatant was not diluted). Isolated DNA was quantified using a Nanodrop 8000 (Thermo Scientific), diluted to 40 ng/4 μL, and stored at -20°C until use in droplet digital PCR for vector genome quantitation. Isolated RNA remained on ice and was quantified using Nanodrop 8000 to ensure relatively equal RNA isolation across samples. RNA was then immediately used for cDNA synthesis with the High-Capacity RNA-to-cDNA™ Kit (Applied Biosystems) according to the kit’s protocol, or frozen at -80°C for later cDNA synthesis. Newly synthesized cDNA was stored at -20°C until use in RT-qPCR for mRNA quantitation.

#### Vector genome quantitation

Vector DNA levels were quantified using duplex TaqMan-based ddPCR assays. The TaqMan reagents used included mTfrc:TaqMan® Copy Number Reference Assay, mouse, Tfrc (Thermo Scientific), and codon-optimized human FOXG1 (opthFOXG1): forward primer (AGCACAGGCCTGACCTTTAT), reverse primer (CATAGGGTGGCTGGGATAGG), and probe (CCCTGCACCACCCAAGGGCC) (Thermo Scientific). ddPCR was performed using the QX200 Droplet Digital PCR System (Bio-Rad). The number of vector genomes per diploid cell (VG/DC) was calculated by dividing the copies/µL obtained with the opthFOXG1 probe set by that obtained with the mTfrc probe set.

#### Human FOXG1 mRNA Quantitation

cDNA was thawed and then added to TaqMan Fast Advance Master mix (Applied Biosystems) as specified in the kit’s protocol with 1:1 ratio of a housekeeper gene (mouse GUSB) to the gene of interest (human FOXG1), TaqMan primer and probe sets. Thermocycler settings were selected according to the TaqMan Fast Advance protocol using QuantStudio 6 (ThermoFisher): Step 1: 50°C, 2 min; Step 2: 95°C, 20 sec.; Step 3: 40 cycles with 95°C, 1 sec. and 60°C, 20 sec.; Step 4: 60°C, 30 sec. The primers and probe for human FOXG1 [Custom TaqMan Gene Expression Assay, VIC (Life Technologies Corporation)] were: forward primer: 5’- AGCACAGGCCTGACCTTTAT, reverse primer: 5’-CATAGGGTGGCTGGGATAGG, probe: 5’-CCCTGCACCACCCAAGGGCC. The primers and probe for mouse GUSB were obtained commercially [Mouse GUSB Assay ID: Mm01197698_m1, FAM (ThermoFisher/ Life Technologies Corporation)].

ΔCq values were calculated by subtracting the housekeeper Cq (mouse GUSB) from the gene of interest Cq (human FOXG1). ΔΔCq values were calculated by subtracting the average ΔCq of the corresponding tissue from the vehicle-treated Q84P group from the ΔCq of each individual sample treated with scAAV9.hSyn1-opthFOXG1. Some samples in these vehicle groups had human FOXG1 Cq’s that were undetermined due to exceeding the Cq cut-off of 40, resulting in incalculable ΔCq’s for these samples, which would skew the average ΔCq per vehicle group higher. Fold change values (2^-ΔΔCq^) were calculated for samples treated with scAAV9.hSyn1-opthFOXG1 relative to the corresponding vehicle-treated Q84P vehicle group (either cortex or hippocampus).

#### Human FOXG1 protein quantitation

Human FOXG1 protein concentrations were quantified by UPLC-MS/MS. Frozen tissues from mouse cortex or hippocampus were thawed, homogenized, denatured for 10 minutes at 100°C, alkylated with iodoacetamide, and digested with trypsin at 37°C. After digestion, the samples were analyzed by UPLC-MS/MS. The human FOXG1 signature peptide (GEPGGGPGELAPVGPDEK) in the mouse cortex and hippocampus tissues were measured. The analysis was accomplished with a SCIEX Triple Quad™ 7500 LC-MS/MS System, which features an IonDrive™ Turbo V source operating in positive electrospray ionization (ESI) mode (SCIEX, Framingham, MA), along with a Shimadzu Nexera XR ultra-high-performance liquid chromatograph (UPLC) system (Shimadzu Scientific Instruments, Japan).

### Immunohistochemical staining for FOXG1

The left brain hemispheres were shipped to Experimental Pathology Laboratories, Inc. (EPL^®^) at Sterling, VA, where they were processed to paraffin embedded blocks, sectioned, and stained with hematoxylin and eosin (H&E), and for FOXG1 by immunohistochemistry. All brain tissues from all study animals were trimmed sagittal and placed in histology cassettes. Tissues in cassettes were processed on an automated tissue processor to dehydrate through increasing ethanol gradients, clear with xylene, and infiltrate tissues with paraffin. Processed tissues were embedded in paraffin, sectioned on a rotary microtome at 4-5 microns, and mounted on glass slides for histochemical and immunohistochemical (IHC) staining. For IHC staining, slides were placed on the Leica BOND RX. The BOND RX instrument deparaffinized slides (Leica, BOND Dewax Solution, Cat# AR9222), unmasked antigens (Leica, BOND Epitope Retrieval Solution 2, Cat# AR9640), and incubated sections in 0.302 µg/mL of recombinant anti-FOXG1 antibody (Abcam, Cat# Ab196868), followed by secondary polymer detection (rabbit) with chromogenic detection and hematoxylin counterstain (Leica, BOND Polymer Refine Detection, Cat# DS9800).

### Neurofilament light chain (NfL) levels

Plasma and CSF samples collected at 12 weeks of age were assessed for NfL using the Quanterix SIMOA assay from Quanterix (Cat# 103400) and the manufacturer’s protocol. CSF and plasma samples were diluted in assay buffer at 1:100 and 1:10, respectively. NfL concentrations were interpolated from a standard curve and corrected for the dilution factor.

### Statistical analysis

Unpaired t-test, ordinary one-way analysis of variance (ANOVA) and two-way mixed effects ANOVAs were used to assess differences in behavioral performance. The Tukey’s or Sidak’s multiple comparisons test was employed for appropriate post-hoc comparisons after the identification of main effects.

## ACKNOWLEDGMENTS

The authors thank the Psychogenics team for in vivo assessments; Richard Miller for assistance in quantifying all DNA and RNA samples as well as preparing DNA samples; Meredith McLerie for overseeing early DNA and RNA isolation workflows and designing the experiments for validating the TaqMan Primer/probe sets for RT-qPCR multiplexing; Dan Zadory and Roza Cheru for immunohistochemistry assistance; Eddy Pichinuk for assistance with coordinating some of the DNA, RNA and histological work; and Eric Smith for assistance with the figures.

---

**Supplemental Figure 1.**
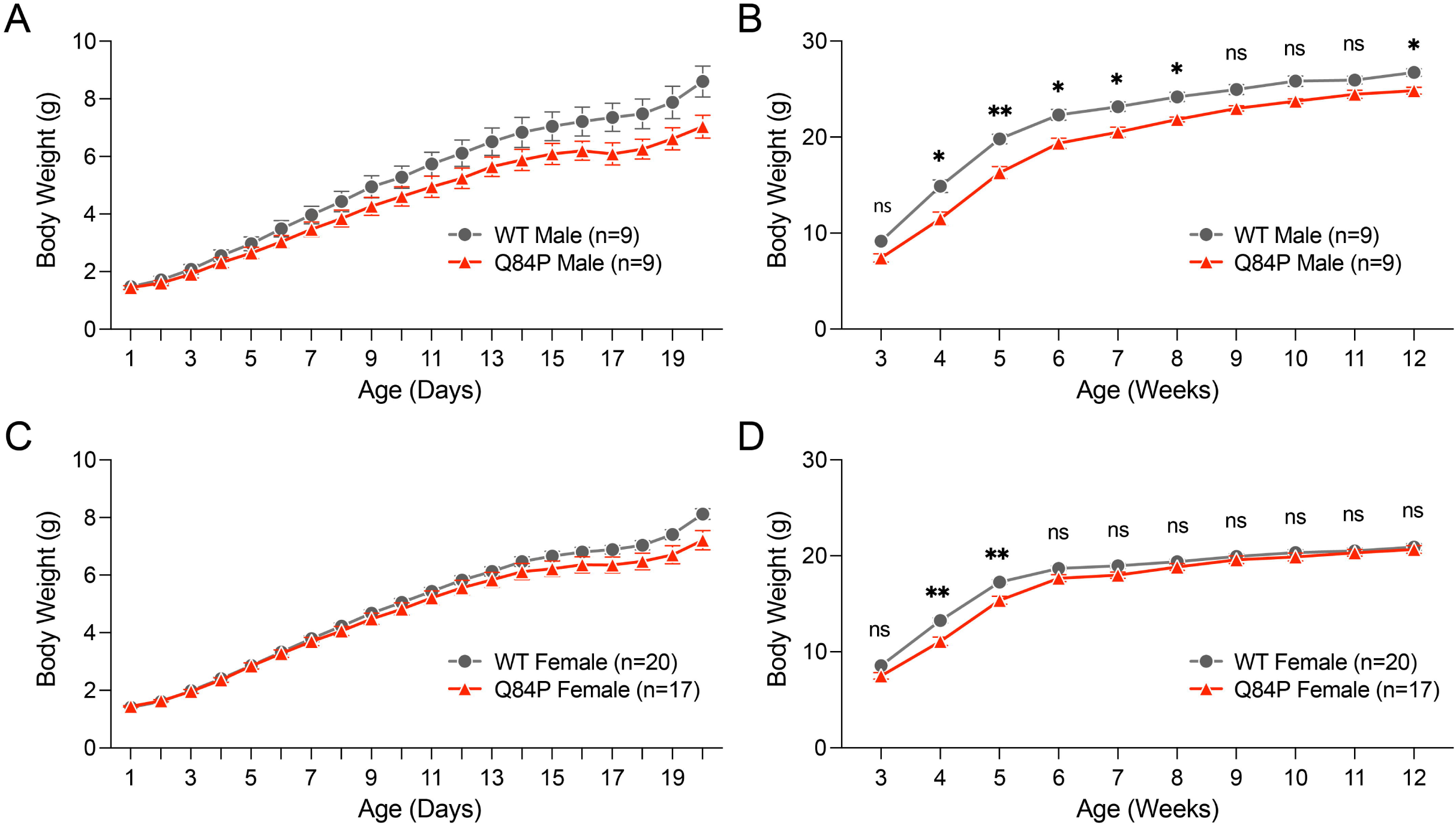
Body weights of male or female Q84P mice analyzed separately from postnatal day 1 to week 12. (**A**) Neonatal male body weights showed no significant differences between genotypes as assessed by one-way ANOVA (F (1, 16) = 2.443, p = 0.1376). (**B**) Adult male body weights showed significant differences between the genotypes as assessed by one-way ANOVA (F (1, 16) = 8.746, p = 0.0093), with significant post-hoc differences at weeks 4 to 8 and 12, as assessed by Tukey’s multiple comparisons test. (**C**) Neonatal female body weights showed no significant differences between genotypes as assessed by one-way ANOVA (F (1, 35) = 1.566, p = 0.2191). (**D**) Adult female body weights showed significant differences between the genotypes as assessed by one-way ANOVA (F (1, 35) = 4.543, p = 0.0401), with significant post-hoc differences at weeks 4 and 5 as assessed by Tukey’s multiple comparisons test. The number of animals per group (n) reflect the total number of animals in each group at the time of enrollment. Means ± standard error of the mean are displayed. ns: not significant. * p < 0.05, ** p < 0.01

**Supplemental Figure 2.**
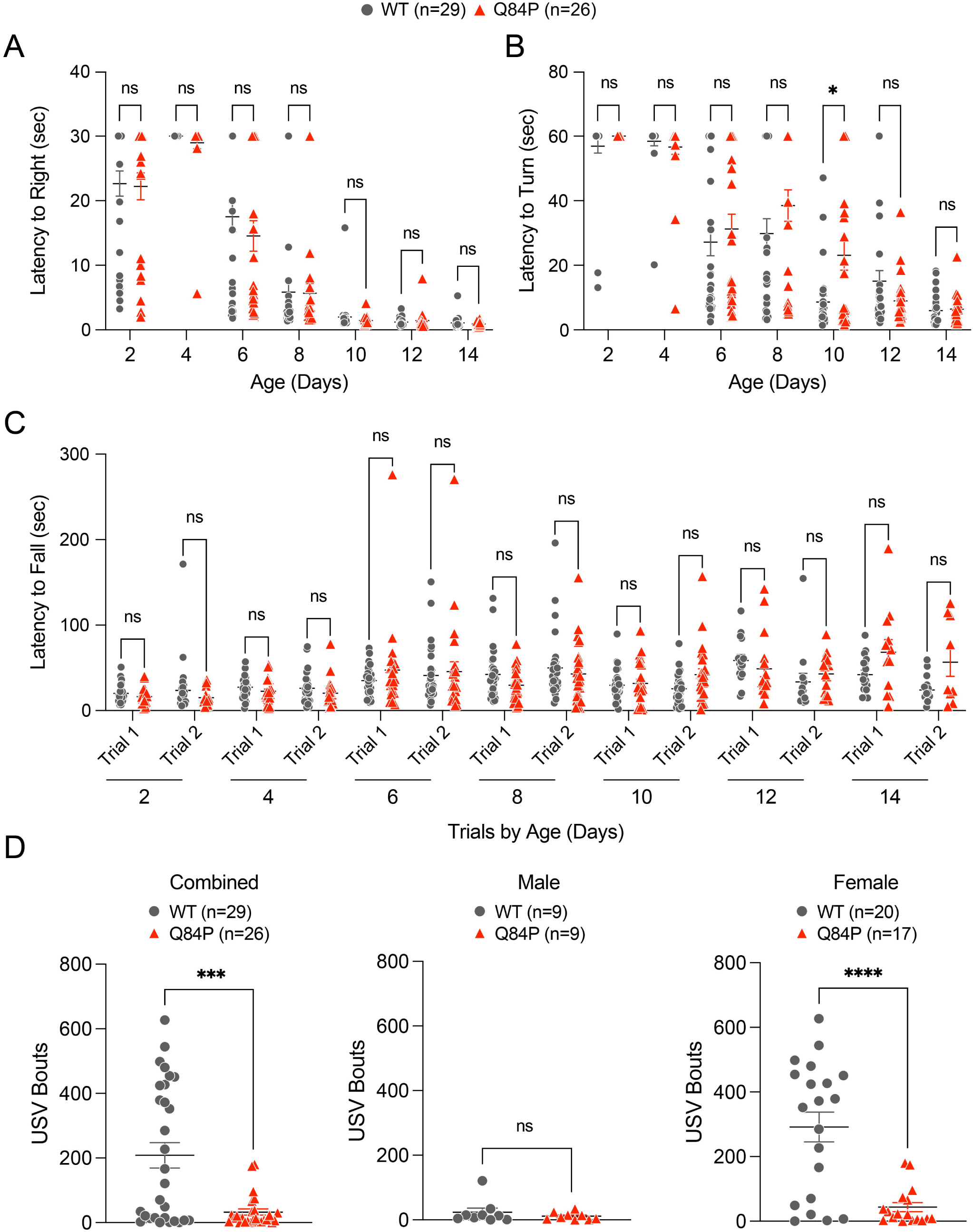
(**A**) Righting reflex assessments in male and female animals combined on postnatal days 2, 4, 6, 8, 10, 12 and 14 showed no significant differences between the genotypes as assessed by one-way ANOVA (F (1, 53) = 0.9778, p = 0.3272). (**B**) Geotaxis assessments in male and female animals combined on postnatal days 2, 4, 6, 8, 10, 12 and 14 showed no significant differences in latency to turn between the genotypes as assessed by one-way ANOVA (F (1, 53) = 3.820, p = 0.0559). (**C**) Tube tests in male and female animals combined on postnatal days 2, 4, 6, 8, 10, 12 and 14 showed no significant differences in latency to fall between the genotypes as assessed by one-way ANOVA (F (1, 53) = 1.763, p = 0.1899). (**D**) Total ultrasonic vocalizations across the 5-minute trial in male and female animals combined on P9 showed significant differences between the genotypes as assessed by one-way ANOVA with Tukey’s multiple comparisons post-hoc test. There were no significant differences in total ultrasonic vocalizations between the genotypes of male animals but there was a significant difference between the genotypes of female animals. Means ± standard error of the mean are displayed. ns: not significant. *** p < 0.001, **** p < 0.0001

**Supplemental Figure 3.**
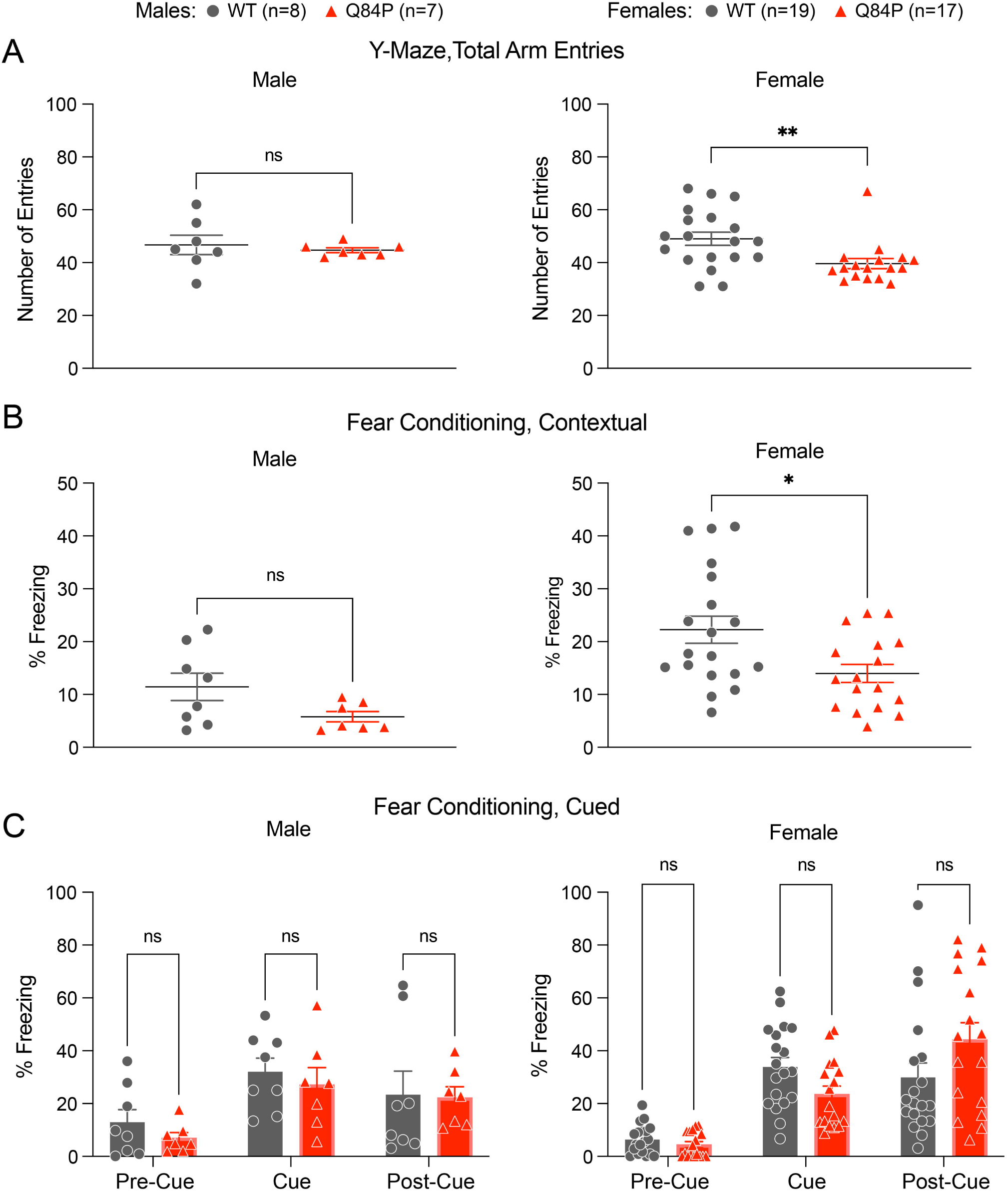
Y-maze total arm entries and contextual fear conditioning are reduced in female but not male Q84P mice compared to WT mice. (**A**) Y-maze total arm entries showed no significant differences between the genotypes in male animals at 9 weeks of age. However, in female mice, there were significant differences in total arm entries between the genotypes. Statistical comparisons were assessed by unpaired t-test. (**B**) Contextual fear conditioning showed no significant difference between the genotypes in male animals at 10 weeks of age. However, in female mice, there were significant differences in contextual percent freezing between the genotypes. Statistical comparisons were assessed by unpaired t-test. (**C**) Cued fear conditioning was not significantly different between the genotypes in either male (F(1, 13) = 0.4084, p = 0.5339) or female (F(1, 34) = 0..02103, p = 0.8856) mice as assessed by two-way ANOVA with Sidak’s multiple comparisons post-hoc test. WT males, n = 8; Q84P males, n = 7; WT females, n = 19; Q84P females, n = 17. Means ± standard error of the mean are displayed. ns: not significant. * p < 0.05, ** p < 0.01

**Supplemental Figure 4.**
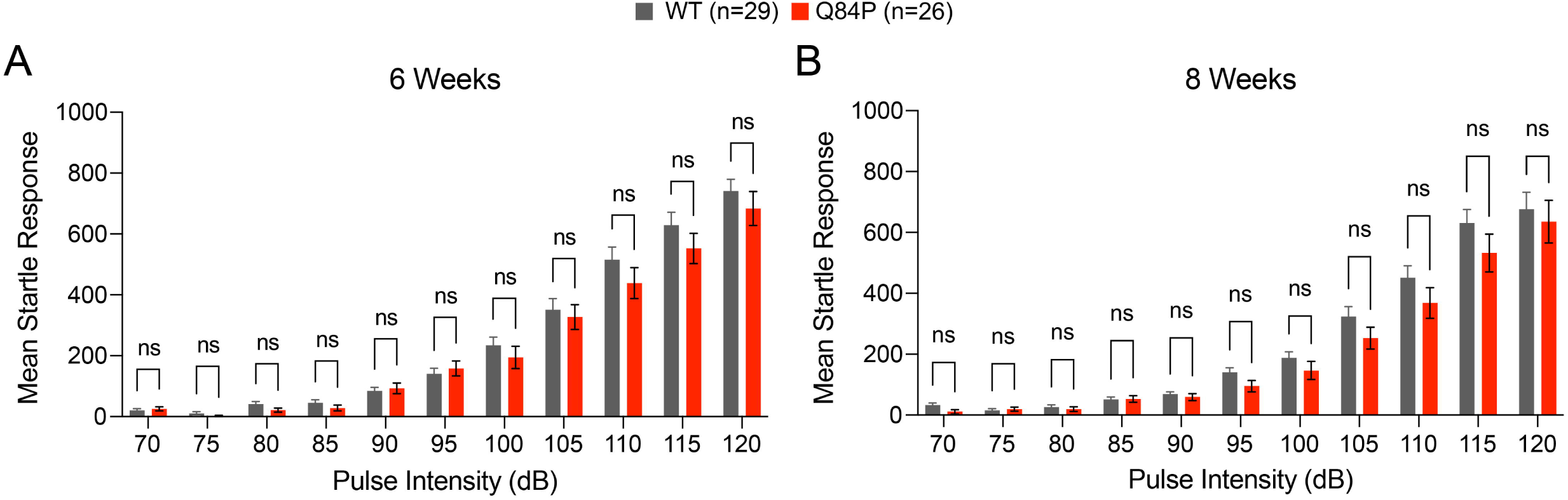
The acoustic startle test showed no significant difference between Q84P and WT mice at 6 weeks of age (F (1, 53) = 0.9232, p = 0.3410) or 8 (F (1, 49) = 1.951, p = 0.1688), as assessed by two-way ANOVA with Sidak’s multiple comparisons post-hoc test. The number of animals per group (n) reflect the total number of combined male and female animals in each group at the time of enrollment. Means ± standard error of the mean are displayed. ns: not significant.

**Supplemental Figure 5.**
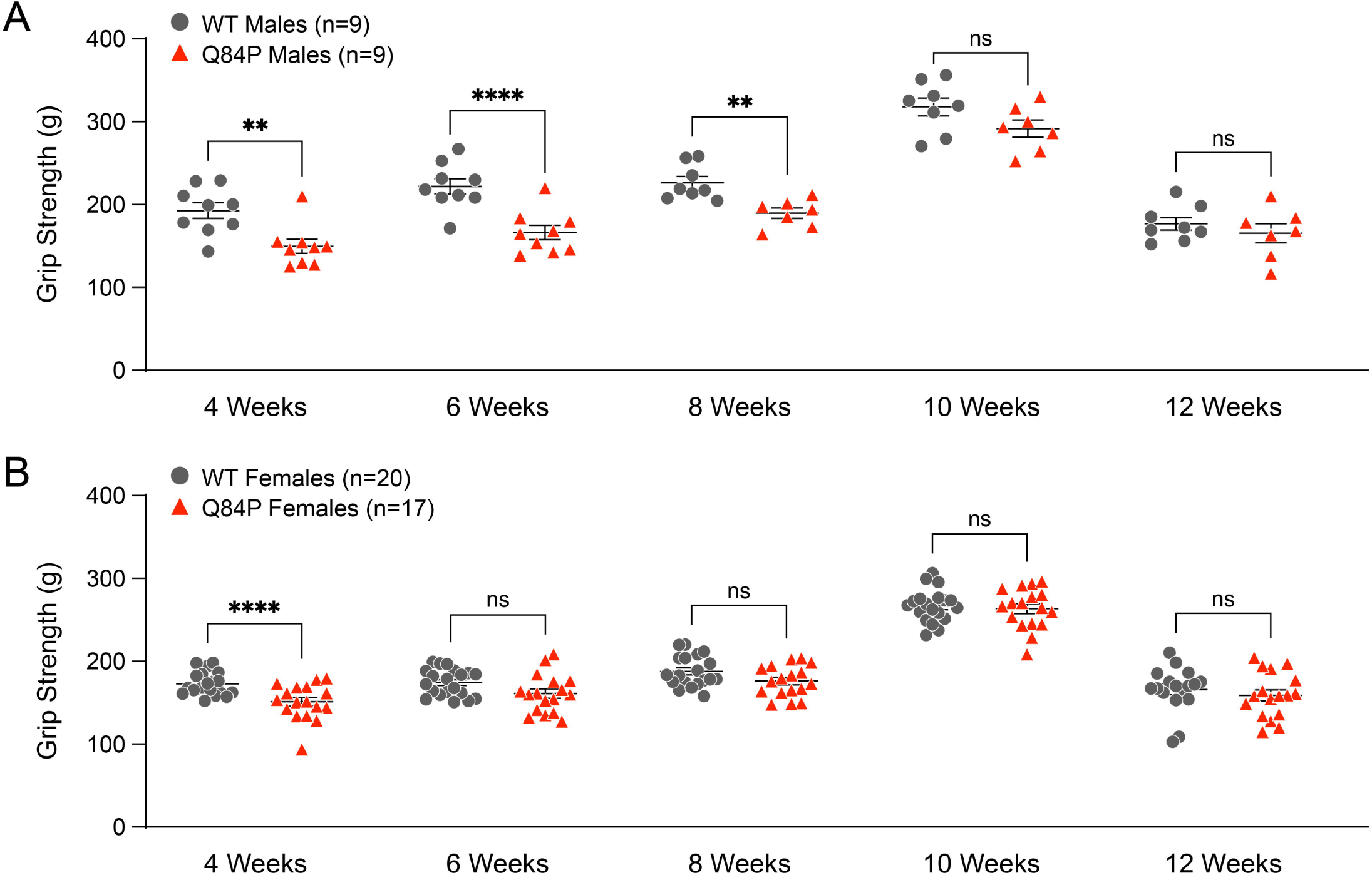
Grip strength is transiently reduced at 4, 6 and 8 weeks of age in male Q84P mice and at only 4 weeks of age in female Q84P mice compared with their WT counterparts. (**A**) Grip strength in males, averaged over 5 trials, showed significant differences between genotypes at 4, 6 and 8 but not 10 or 12 weeks of age. (**B**) Grip strength in females, averaged over 5 trials, showed significant differences between genotypes at 4 but not 6, 8, 10 or 12 weeks of age. Statistical comparisons were assessed by unpaired t-test. The number of animals per group (n) reflect the total number of animals in each group at the time of enrollment. Means ± standard error of the mean are displayed. ns: not significant. ** p < 0.01, **** p < 0.0001

**Supplemental Figure 6.**
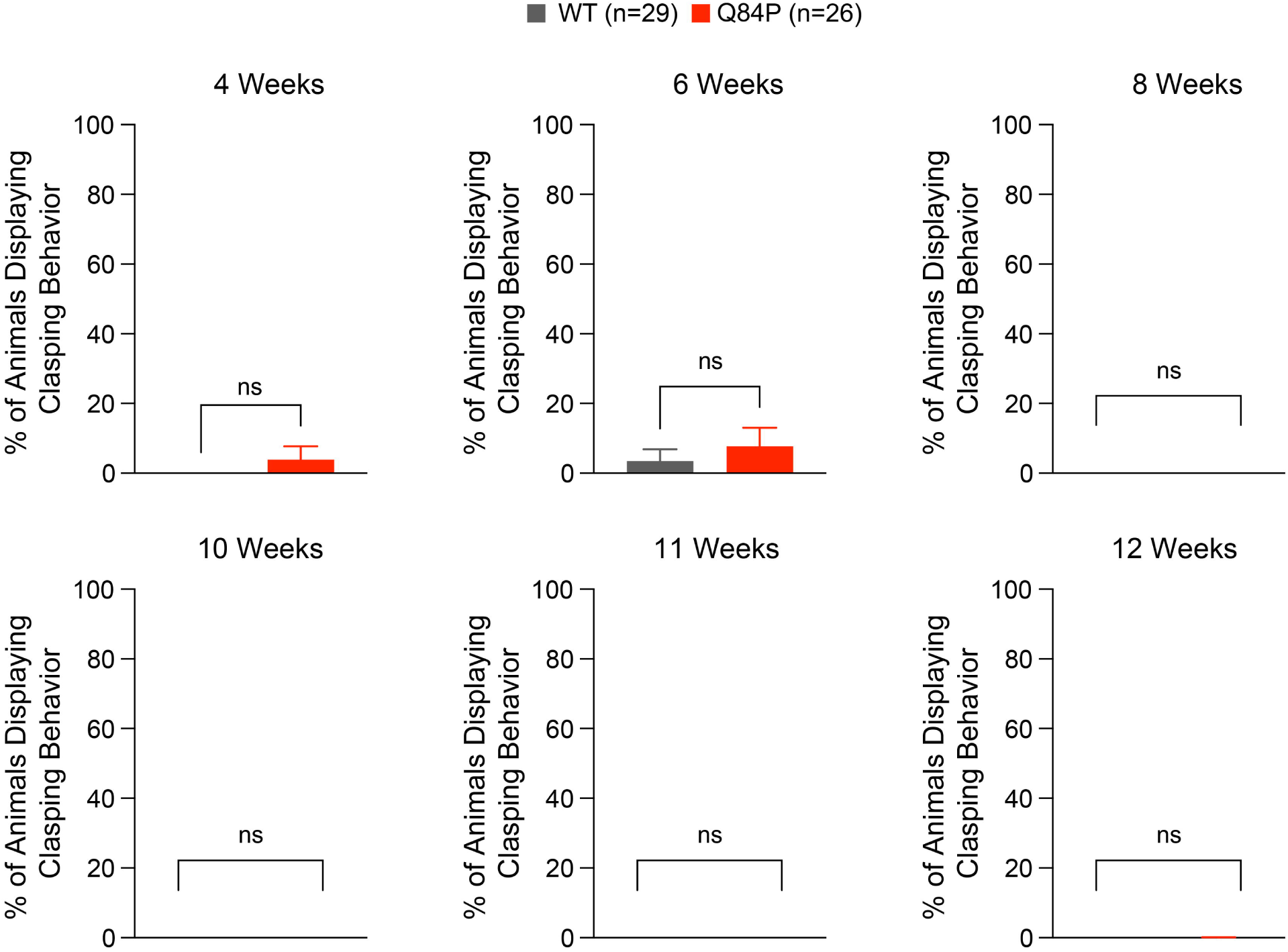
Clasping behavior showed no significant difference between Q84P and WT mice at any age examined. There were no significant differences in the percentage of animals exhibiting clasping at 4 or 6 weeks of age. At 8, 10, 11 and 12 weeks of age, there was no clasping observed in either Q84P or WT mice. Statistical comparisons were assessed by unpaired t-test. The number of animals per group (n) reflect the total number of combined male and female animals in each group at the time of enrollment. Means ± standard error of the mean are displayed. ns: not significant.

**Supplemental Figure 7.**
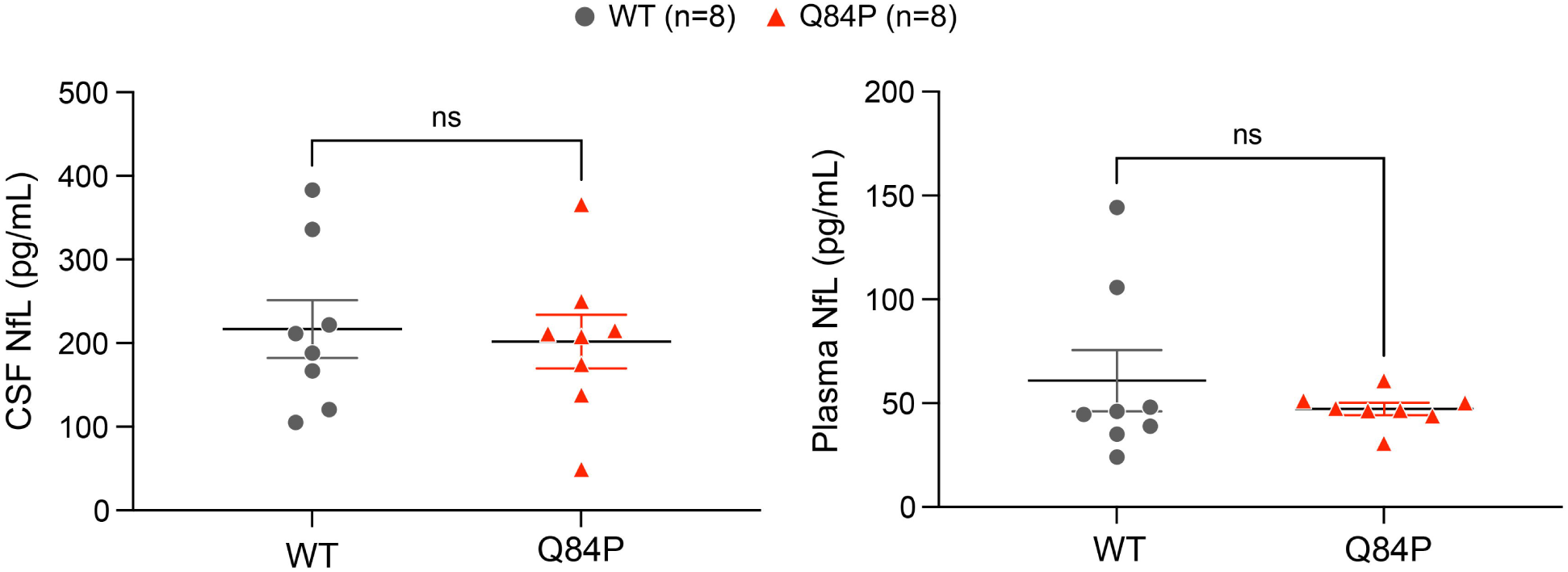
NfL protein levels in CSF (**A**) and plasma (**B**) showed no significant differences between Q84P and WT mice at 12 weeks of age. Statistical comparisons were assessed by unpaired t-test. Means ± standard error of the mean are displayed. ns: not significant.

**Supplemental Figure 8.**
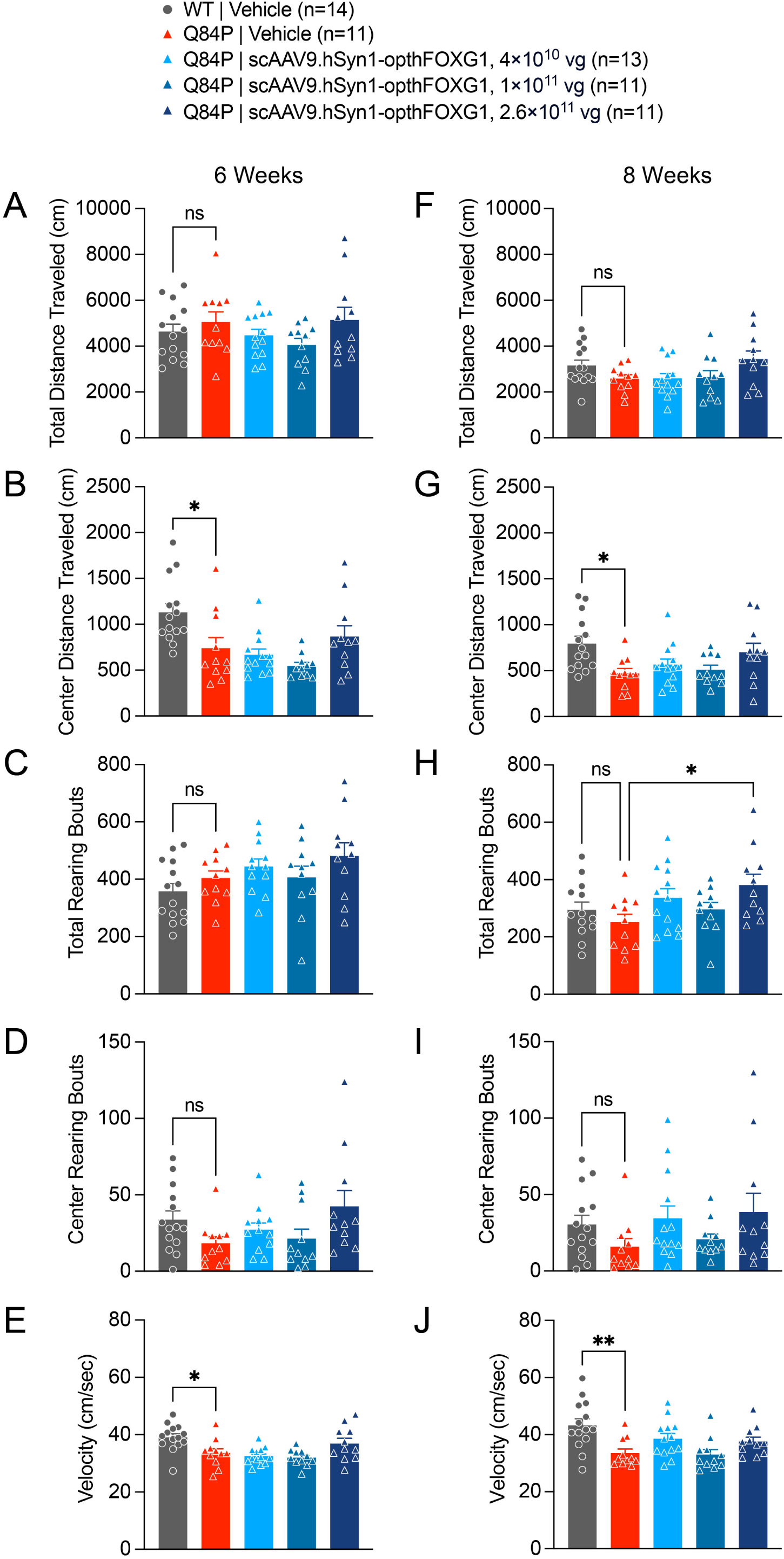
Open field center distance travelled and velocity were significantly reduced in 6- and 8-week old Q84P compared to WT mice but not normalized by scAAV9.hSyn1-opthFOXG1 treatment at any ICV dose tested (4 × 10^10^, 1 × 10^11^ or 2.6x10^11^ vg per mouse). Decreases in other endpoints such as total distance travelled, and total and center rearing bouts did not reach statistical significance in vehicle-treated Q84P vs WT mice. Open field total (**A, F**) and center (**B, G**) distance travelled, total (**C, H**) and center (**D, I**) rearing frequency, and velocity (**E, J**) are shown at 6 (**A-E**) and 8 (**F-J**) weeks of age. The only effect of ICV treatment with scAAV9.hSyn1-opthFOXG1 was a significant increase in the total rearing bouts at 8 weeks of age with a dose of 2.6x10^11^ vg per mouse compared to vehicle. At each age, one-way ANOVA with Tukey’s multiple comparisons test was used to assess statistical significance. Means ± standard error of the mean are displayed. ns: not significant. * p < 0.05, ** p < 0.01

**Supplemental Figure 9.**
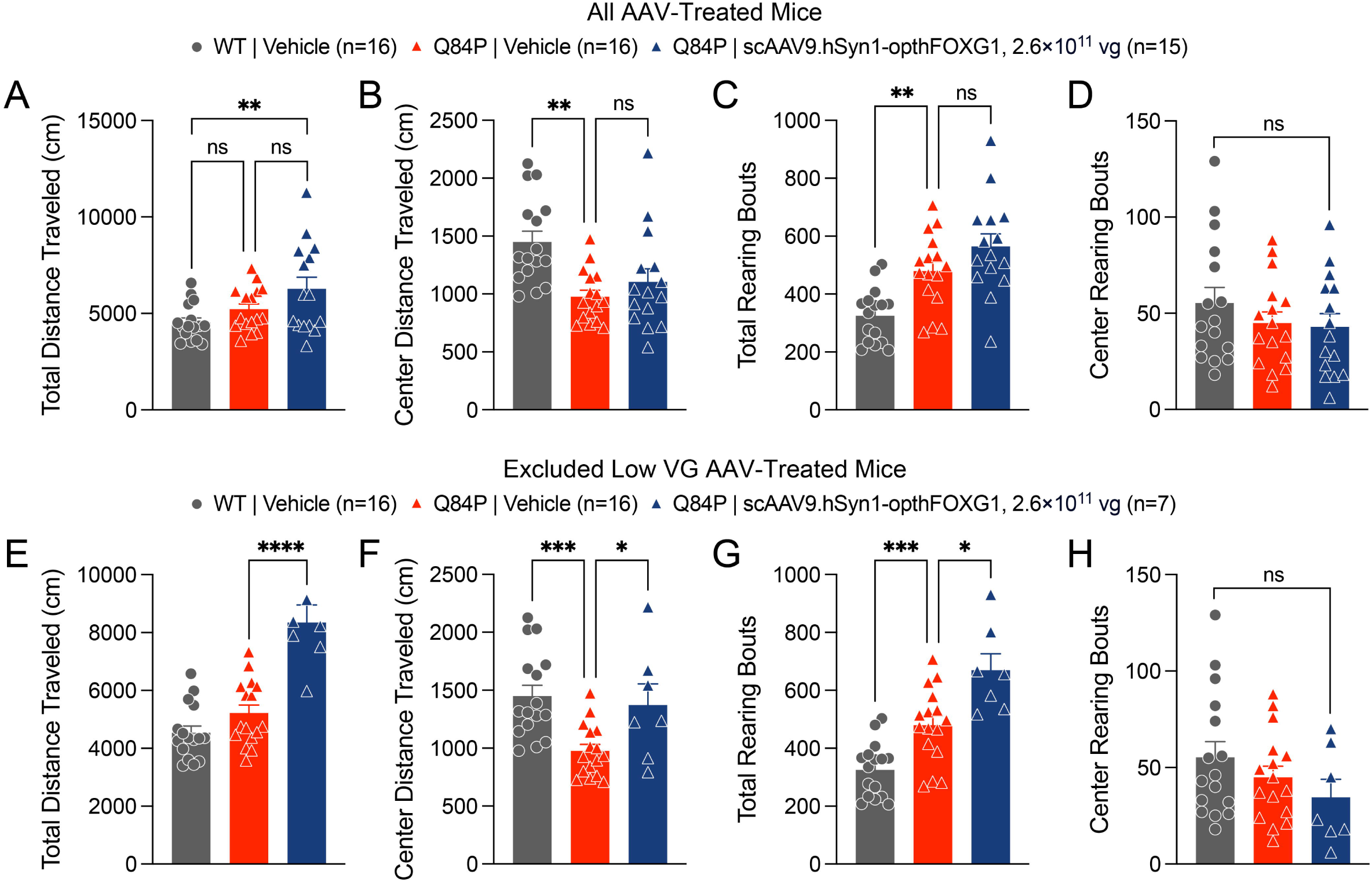
Open field total and center distance travelled and total and center rearing frequency in 11-week old vehicle-treated WT and Q84P mice and Q84P mice treated with scAAV9.hSyn1-opthFOXG1 at an ICV dose of 2.6 × 10^11^ vg per mouse. Open field total (**A, E**) and center (**B, F**) distance travelled, and total (**C, G**) and center (**D, H**) rearing frequency are shown for all animals (**A-D**) and after exclusion of eight Q84P mice in the scAAV9.hSyn1-opthFOXG1 group that exhibited less than one-tenth of the expected levels of vector genomes in hippocampus and cortex (**E-H**). When all animals were assessed by one-way ANOVA with Tukey’s multiple comparisons test (**A-D**), there were significant differences in total distance travelled among the treatment groups (F (2, 44) = 4.845, p = 0.0125), in center distance travelled among the treatment groups (F (2, 44) = 7.607, p = 0.0015), and in total rearing frequency among the treatment groups (F (2, 44) = 12.98, p < 0.0001), but not in center rearing frequency among the treatment groups. When the eight Q84P mice treated with scAAV9.hSyn1-opthFOXG1 that had less than one-tenth of expected vector genome levels in cortex and hippocampus tissues were excluded (**E-H**), there was a significant increase in center distance travelled and total rearing frequency as well as total distance travelled after scAAV9.hSyn1-opthFOXG1 versus vehicle treatment in Q84P mice but no change in center rearing frequency. Means ± standard error of the mean are displayed. ns: not significant. * p < 0.05, ** p < 0.01, *** p < 0.001, **** p < 0.0001

**Supplemental Figure 10.**
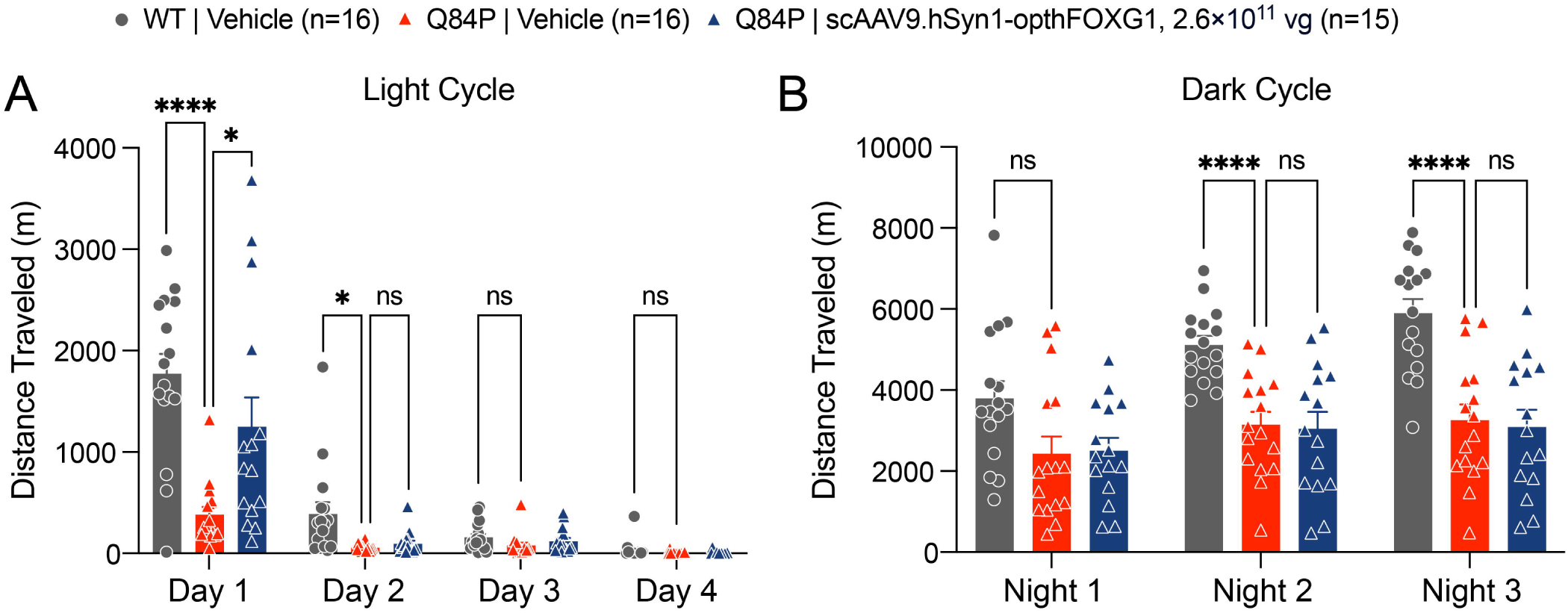
Running wheel total distance travelled during the light and dark cycles at 12 weeks of age in all mice, including those treated with scAAV9.hSyn1-opthFOXG1 that had less than one-tenth of expected vector genome levels in cortex and hippocampus tissues. (**A**) The total distance travelled during the animals’ light cycle (from 7AM to 7PM) was assessed on 4 days and showed significant differences between the treatment groups, as assessed by two-way ANOVA (F (2, 44) = 14.75, p < 0.0001). Specifically, there were significant reductions in vehicle-treated Q84P compared with WT mice on Day 1 and Day 2, as assessed with Tukey’s multiple comparisons post-hoc test, with a significant partial normalization on Day 1 in scAAV9.hSyn1-opthFOXG1-treated Q84P mice. (**B**) The total distance travelled during the animals’ dark cycle (from 7PM to 7AM) was assessed on 3 nights and showed significant differences in the total distance travelled between the treatment groups, as assessed by two-way ANOVA (F (2, 44) = 13.03, p < 0.0001). Specifically, there were significant reductions in Q84P mice compared with WT mice treated with vehicle on Night 2 and Night 3, as assessed with Tukey’s multiple comparisons post-hoc test. There was no significant effect of treatment with scAAV9.hSyn1-opthFOXG1 on either night. Means ± standard error of the mean are displayed. ns: not significant * p < 0.05, **** p< 0.0001

